# The role of structural pleiotropy and regulatory evolution in the retention of heteromers of paralogs

**DOI:** 10.1101/564401

**Authors:** Axelle Marchant, Angel F. Cisneros, Alexandre K Dubé, Isabelle Gagnon-Arsenault, Diana Ascencio, Honey A. Jain, Simon Aubé, Chris Eberlein, Daniel Evans-Yamamoto, Nozomu Yachie, Christian R Landry

## Abstract

Gene duplication is a driver of the evolution of new functions. The duplication of genes encoding homomeric proteins leads to the formation of homomers and heteromers of paralogs, creating new complexes after a single duplication event. The loss of these heteromers may be required for the two paralogs to evolve independent functions. Using yeast as a model, we find that heteromerization is frequent among duplicated homomers and correlates with functional similarity between paralogs. Using *in silico* evolution, we show that for homomers and heteromers sharing binding interfaces, mutations in one paralog can have structural pleiotropic effects on both interactions, resulting in highly correlated responses of the complexes to selection. Therefore, heteromerization could be preserved indirectly due to selection for the maintenance of homomers, thus slowing down functional divergence between paralogs. We suggest that paralogs can overcome the obstacle of structural pleiotropy by regulatory evolution at the transcriptional and post-translational levels.

## Introduction

Proteins assemble into molecular complexes that perform and regulate structural, metabolic and signalling functions (Janin et al., 2008; Marsh and Teichmann, 2015; Pandey et al., 2017; Scott and Pawson, 2009; Vidal et al., 2011; Wan et al., 2015). The assembly of complexes is necessary for protein function and thus constrains the sequence space available for protein evolution. One direct consequence of protein-protein interactions (PPIs) is that a mutation in a given gene can have pleiotropic effects on other genes’ functions through physical associations. Therefore, to understand how genes and cellular systems evolve, we need to consider physical interactions as part of the environmental factors shaping a gene’s evolutionary trajectory (Landry et al., 2013; Levy et al., 2012).

A context in which PPIs and pleiotropy may be particularly important is during the evolution of new genes after duplication events (Amoutzias et al., 2008; Baker et al., 2013; Diss et al., 2017; Kaltenegger and Ober, 2015). The molecular environment of a protein in this context includes its paralog if the duplicates derived from an ancestral gene encoding a self-interacting protein (homomer) (Figure 1). In this case, mutations in one paralog could have functional consequences for the other copy because the duplication of a homomeric protein (HM) leads not only to the formation of two new homomers (HMs) but also to a new heteromer (HET) (Figure 1) (Pereira-Leal et al., 2007; Wagner, 2003). We refer to these complexes as homomers and heteromers of paralogs.

**Figure 1:**
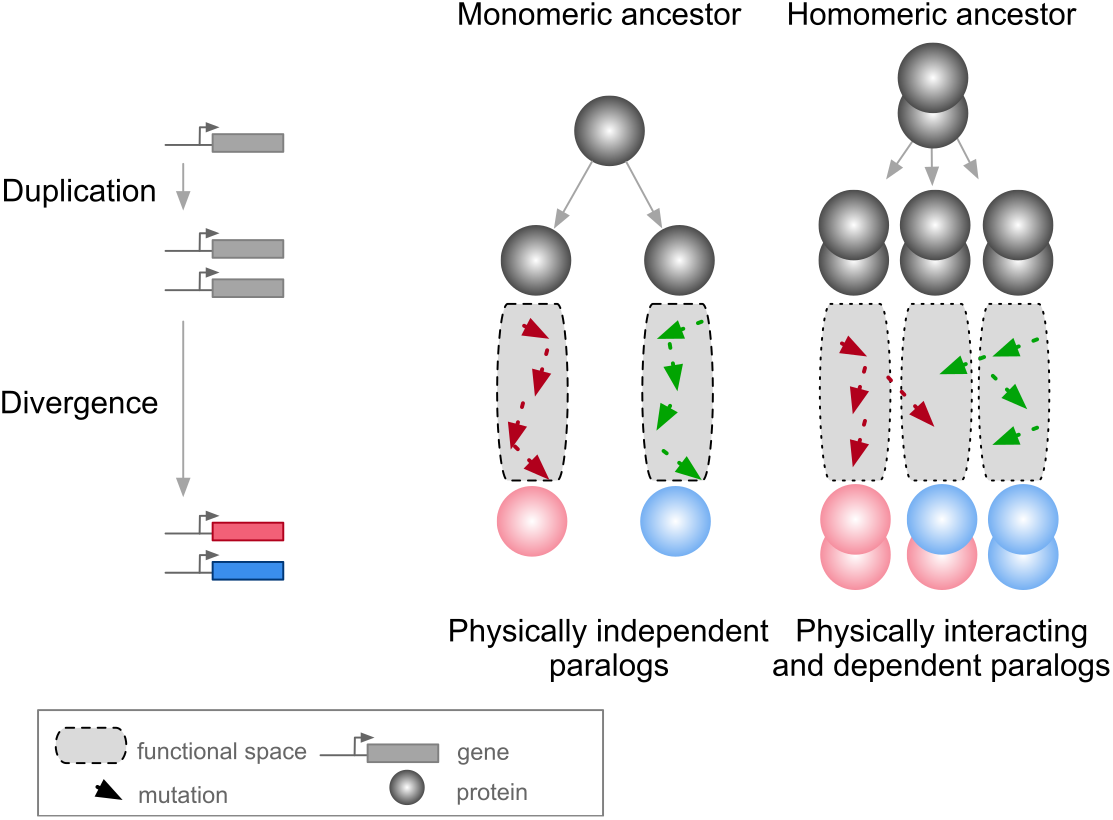
Mutations in paralogous proteins originating from an ancestral homomer are likely to have pleiotropic effects on each other’s function due to their physical association. Gene duplication leads to physically interacting paralogs when they derive from an ancestral homomeric protein. The evolutionary fates of the physically associated paralogs tend to be interdependent because mutations in one gene can impact on the function of the other copy through heteromerization.

Paralogs originating from HMs are physically associated as HETs when they arise. Subsequent evolution can lead to the maintenance or the loss of these HETs. Consequently, paralogs that maintained the ability to form HETs have often evolved new functional relationships (Amoutzias et al., 2008; Baker et al., 2013; Kaltenegger and Ober, 2015). Examples include a paralog degenerating and becoming a repressor of the other copy (Bridgham et al., 2008), and pairs of paralogs that split the functions of the ancestral HM between one of the HMs and the HET (Baker et al., 2013), that cross-stabilize and thus need each other to perform their function (Diss et al., 2017) or that evolved a new function together as a HET (Boncoeur et al., 2012). However, there are also paralogs that do form HMs but that have lost the ability to form HETs through evolution. Among these are duplicated histidine kinases (Ashenberg et al., 2011) and many heat-shock proteins (Hochberg et al., 2018). For the majority of HETs, we do not know what novel functions, if any, contribute to their maintenance.

Therefore, one important question to examine is: what are the evolutionary forces at work for the maintenance or the disruption of HETs arising from HMs? Previous studies suggest that if a paralog pair maintains its ability to form HMs, it is very likely to maintain the HET complex as well (Pereira-Leal et al., 2007). For instance, Lukatsky et al. (Lukatsky et al., 2007) showed that proteins tend to intrinsically interact with themselves and that negative selection may be needed to disrupt HMs. Given this, and since nascent paralogs are identical just after duplication, they would tend to maintain a high propensity to assemble with each other. Hence, the two paralogs would form both HMs and HETs until mutations that destabilize one or the other specifically accumulate (Ashenberg et al., 2011; Hochberg et al., 2018). In addition, the rate at which the HET is lost may depend on the combined effects of mutations in the different subunits since epistasis may cause mutations together to be more or less disruptive for the HET than for the HMs (Diss and Lehner, 2018; Starr and Thornton, 2016). Here, we hypothesize that the association of paralogs forming HETs acts as a constraint that may slow the functional divergence of paralogs by keeping gene products physically associated.

Previous studies have shown that HMs are enriched in eukaryotic PPI networks (Lynch, 2012; Pereira-Leal et al., 2007). However, the extent to which paralogs interact with each other has not been comprehensively quantified in any species. We therefore examine the physical assembly of paralogous proteins (HETs) exhaustively in a eukaryotic interactome by integrating data from the literature and by performing a large-scale PPI screening experiment. Second, using functional data analysis, we examine the consequences of losing HET formation for HM forming paralogs. We perform *in silico* evolution experiments to examine whether the molecular pleiotropy of mutations, caused by shared binding interfaces between HM and HET complexes, could contribute to maintain interactions between paralogs originating from ancestral HMs. We show that selection to maintain HMs alone may be sufficient to prevent the loss of HETs. Finally, we find that regulatory evolution, either at the level of gene transcription or protein localization, may relieve the pleiotropic constraints maintaining the interaction of paralogous proteins.

## Results

### Homomers among singletons and paralogs in the yeast PPI network

We first examined the extent of homomerization across the yeast proteome (see dataset in methods and the supplementary text) for two classes of paralogs, those that are small-scale duplicates (SSDs) and those that are whole-genome duplicates (WGDs). We considered these two sets separately because they may have been retained through different mechanisms (see below). The dataset for this analysis, which includes previously reported PPIs and novel DHFR Protein-fragment Complementation Assay experiments (referred to as PCA, see methods and supplementary text), covers 2521 singletons, 2547 SSDs, 866 WGDs and 136 genes that are both SSDs and WGDs (henceforth referred to as 2D) (Tables S1 and S2). We find that among the 6070 tested yeast proteins, 1944 (32%) form HMs, which agrees with previous estimates from crystal structures (Lynch, 2012). The proportion of HMs among singletons (n = 630, 25%) is lower than for all duplicates: SSDs (n = 980, 38%, p-value < 2.0e-16), WGDs (n = 283, 33%, p-value = 1.6e-05) and 2D (n = 51, 38%, p-value = 1.7e-03) (Figure 2. A, Tables S1 and S2).

**Figure 2:**
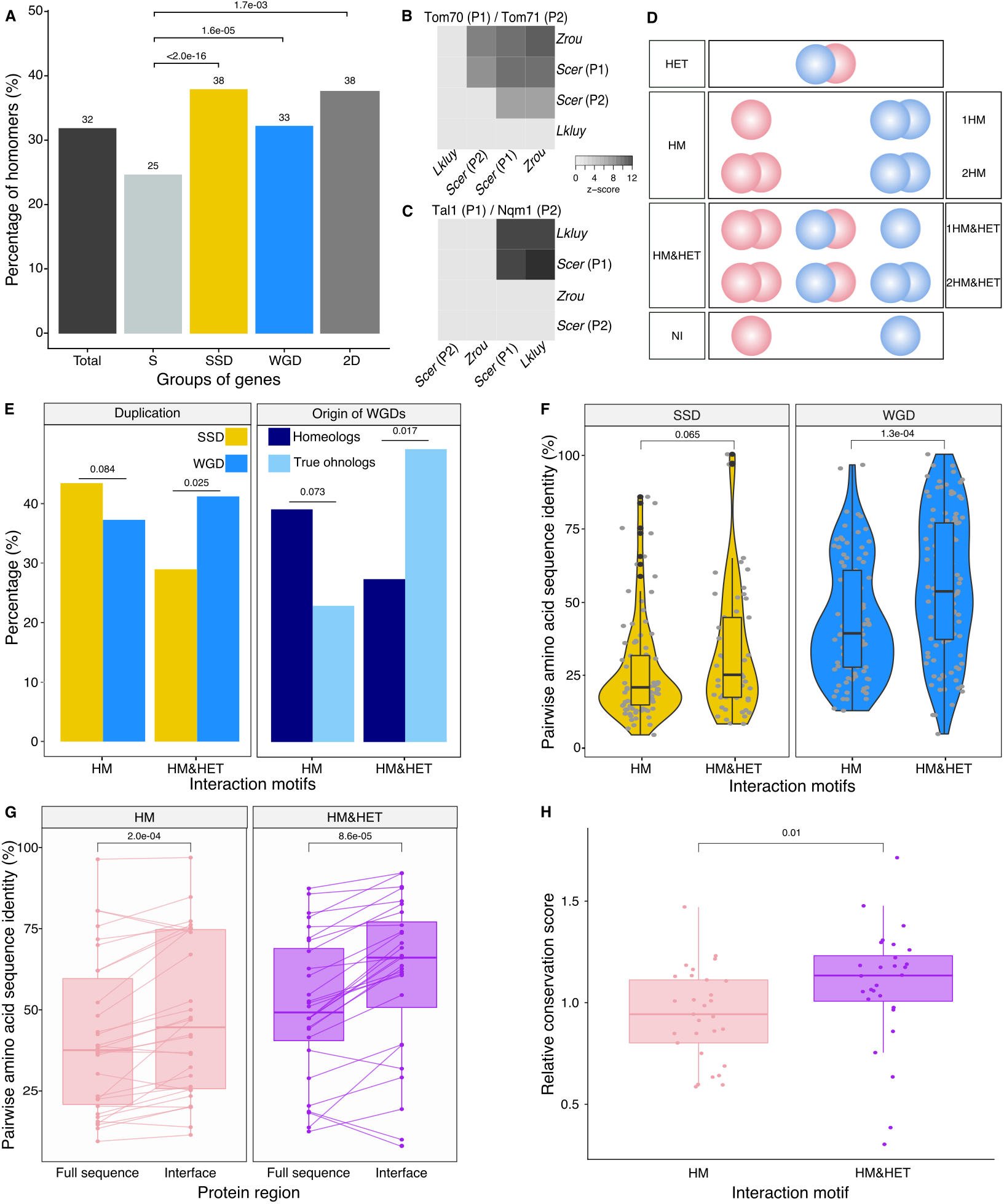
Homomers and heteromers of paralogs are frequent in the yeast protein interaction network. **(A)** The percentage of homomeric proteins in *S. cerevisiae* varies among singletons (S, n = 2521 tested), small-scale duplicates (SSDs, n = 2547 tested), whole-genome duplicates (WGDs, n = 866 tested) and genes duplicated by the two types of duplication (2D, n = 136 tested) (global Chi-square test: p-value < 2.2e-16). Each category is compared with the singletons using a Fisher’s exact test. P-values are reported on the graph. (**B and C**) Interactions between *S. cerevisiae* paralogs and pre-whole-genome duplication orthologs using DHFR PCA. The gray tone shows the PCA signal intensity converted to z-scores. Experiments are performed in *S. cerevisiae*. Interactions are tested among: (**B**) *S. cerevisiae* (*Scer*) paralogs Tom70 (P1) and Tom71 (P2) and their orthologs in *Lachancea kluyveri* (*Lkluy*, SAKL0E10956g) and in *Zygosaccharomyces rouxii* (*Zrou*, ZYRO0G06512g) and (**C**) *S. cerevisiae* paralogs Tal1 (P1) and Nqm1 (P2) and their orthologs in *L. kluyverì* (*Lkluy*, SAKL0B04642g) and in *Z. rouxii* (*Zrou*, ZYRO0A12914g). (**D**) Paralogs show six interaction motifs that we grouped in four categories according to their patterns. HET pairs show heteromers only. HM pairs show at least one homomer (one for 1HM or two for 2HM). HM&HET pairs show at least one homomer (one for 1 HM&HET or two for 2HM&HET) and the heteromer. NI (non-interacting) pairs show no interaction. We focused our analysis on pairs derived from an ancestral HM, which we assume are pairs showing the HM and HM&HET motifs. (**E**) Percentage of HM and HM&HET among SSDs (202 pairs considered, yellow) and WGDs (260 pairs considered, blue) (left panel), homeologs that originated from inter-species hybridization (47 pairs annotated and considered, dark blue) (right panel) and true ohnologs from the whole-genome duplication (82 pairs annotated and considered, light blue). P-values are from Fisher’s exact tests. (**F**) Percentage of pairwise amino acid sequence identity between paralogs for HM and HM&HET motifs for SSDs and WGDs. P-values are from Wilcoxon tests. (**G**) Pairwise amino acid sequence identity for the full sequences of paralogs and their binding interfaces for the two motifs HM and HM&HET. P-values are from paired Wilcoxon tests. (**H**) Relative conservation scores for the two motifs of paralogs. Conservation scores are the percentage of sequence identity at the binding interface divided by the percentage of sequence identity outside the interface. Data shown include 30 interfaces for the HM group and 28 interfaces for the HM&HET group (22 homomers and 3 heterodimers of paralogs) (Table S13). P-value is from a Wilcoxon test. Figure 2 - figure supplements 1 to 8.

Although a large number of PPIs have been previously reported in *S. cerevisiae*, it is possible that the frequency of HMs is slightly underestimated because they were not systematically and comprehensively tested (see methods). Another reason could be that some interactions could not be detected due to low expression levels. We measured mRNA abundance in cells grown in PCA conditions and used available yeast protein abundance data (Wang et al., 2012) to test this possibility (Tables S3, S4, S5 and S6). As previously observed (Celaj et al., 2017; Freschi et al., 2013), we found a correlation between PCA signal from our experiments and expression level, both at the level of mRNA and protein abundance (Spearman r = 0.33, p-value = 3.5e-13 and Spearman r = 0.46, p-value < 2.2e-16 respectively). When focusing only on HMs previously reported, we also observed both correlations (Spearman r = 0.37, p-value = 3.9e-08 and Spearman r = 0.38, p-value = 6.0e-08 respectively). The association between PCA signal and expression translates into a roughly two-fold increase in the probability of HM detection when mRNA levels change by one order of magnitude (Figure 2–figure supplement 3. A). We also generally detected stronger PCA signal for the HM of the most expressed paralog of a pair, confirming the effect of expression on our ability to detect PPIs (Figure 2–figure supplement 3. B). Finally, we found that HMs reported in the literature but not detected by PCA have on average lower expression levels (Figure 2–figure supplement 3. B-C). We therefore conclude that some HMs (and also HETs) remain undetected because of low expression levels.

The overrepresentation of HMs among duplicates was initially observed for human paralogs (Pérez-Bercoff et al., 2010). One potential mechanism to explain this finding is that homomeric proteins are more likely to be maintained as pairs after duplication because they might become dependent on each other for their stability that is enhanced through the formation of HET (Diss et al., 2017). Another explanation is that proteins forming HMs could be expressed at higher levels and therefore, easier to detect, as shown above. High expression could also itself increase the long term probability of genes to persist after duplication (Gout et al., 2010; Gout and Lynch, 2015). We indeed observed that both SSDs and WGDs are more expressed than singletons at the mRNA and protein levels, with WGDs being more expressed than SSDs at the mRNA level (Figure 2–figure supplement 4. A-B). However, expression level (and thus PPI detectability) does not explain completely the enrichment of HMs among duplicated proteins. Both factors, expression and duplication, have significant effects on the probability of proteins to form HMs (Table S7. A). It is therefore likely that the overrepresentation of HMs among paralogs is linked to their higher expression but other factors are also involved.

### Paralogous heteromers frequently derive from ancestral homomers

The model presented in Figure 1 assumes that the ancestral protein leading to HET formed a HM before duplication. Under the principle of parsimony, we can assume that when at least one paralog forms a HM, the ancestral protein was also a HM. This was shown to be true in general by (Diss et al., 2017) that compared yeast WGDs to their orthologs from *Schizosaccharomyces pombe*. To further support this observation, we used PCA to test for HM formation for orthologs from species that diverged prior to the whole-genome duplication event (*Lachancea kluyveri* and *Zygosaccharomyces rouxii*). We looked at the mitochondrial translocon complex and at the transaldolase, which both show HETs (see methods). We confirm that when one HM was observed in *S. cerevisiae*, at least one ortholog from pre-whole-genome duplication species formed a HM (Figure 2. B-C). We also detected interactions between orthologs, suggesting that ability to interact has been preserved despite the millions of years of evolution separating these species. The absence of interactions for some of these orthologous proteins may be due to the incompatibility of their expression in *S. cerevisiae*.

We then focused on HMs and HETs for 202 pairs of small-scale duplicates (SSDs) and 260 pairs of whole-genome duplicates (WGDs). It is a reduced dataset compared to the previous section because we needed to consider only pairs for which there was no missing PPI data (see methods). We combined public data with our own PCA experimental data on 86 SSDs and 149 WGDs (see supplementary text, Figure 2–figure supplement 1 and 2). Overall, the data represents a total of 462 pairs of paralogs (202 SSDs and 260 WGDs) covering 53% of the SSDs and 50% of the WGDs (Tables S3 and S4). This dataset covers 493 binary interactions of paralogs with themselves (HMs) and 214 interactions with their sister copy (HET).

We classified paralog pairs into four classes according to whether they show only the HET (HET, 10%), at least one HM but no HET (HM, 39%), at least one of the HM and the HET (HM&HET, 37%) or no interaction (NI, 15%) (Figure 2. D, supplementary text). Overall, most pairs forming HETs also form at least one HM (79%, Table S3). For the rest of the study, we focused our analysis and comparisons on HM and HM&HET pairs because they most likely derive from an ancestral HM. Previous observations showed that paralogs are enriched in protein complexes comprising more than two distinct subunits, partly because complexes evolved by the initial establishment of self-interactions followed by duplication of homomeric proteins (Musso et al., 2007; Pereira-Leal et al., 2007). However, we find that the majority of HM&HETs could be simple oligomers of paralogs that do not involve other proteins and are thus not part of large complexes. Only 70 (41%) of the 169 cases of HM&HET are in complexes with more than two distinct subunits among a set of 5,535 complexes reported in databases (see methods).

We observed that the correlation between HM and HET formation is affected by whether paralogs derived from SSD or WGD (Figure 2. E). WGDs tend to form HETs more often when they form at least one HM, resulting in a larger proportion of HM&HET motif than SSDs. We hypothesize that since SSDs have appeared at different evolutionary times, many of them could be older than WGDs, which could be accompanied by a loss of interactions between paralogs. Indeed, we observed that the distribution of sequence divergence shows lower identity for SSDs than for WGDs, suggesting the presence of ancient duplicates that predate the whole-genome duplication (Figure 2–figure supplement 5. A). Higher protein sequence divergence could lead to the loss of HET complexes because it increases the probability of divergence at the binding interface. We indeed found that among SSDs, those forming HM&HET tend to show a marginally higher overall sequence identity (p=0.065, Figure 2. F, Figure 2–figure supplement 5. B and C). We also observed a significantly higher sequence identity for WGD pairs forming HM&HET, albeit with a wider distribution (Figure 2. F, Figure 2–figure supplement B C). This wider distribution at least partly derives from the mixed origin of WGDs (Figure 2-figure supplement 5). Recently, Marcet-Houben and Gabaldón (Marcet-Houben and Gabaldón, 2015; Wolfe, 2015) showed that WGDs likely have two distinct origins: actual duplication (generating true ohnologs) and hybridization between species (generating homeologs). For pairs whose ancestral state was a HM, we observed that true ohnologs have a tendency to form HET more frequently than homeologs (Figure 2. E). Because homeologs had already diverged before the hybridization event, they are older than ohnologs, as shown by their lower pairwise sequence identity (Figure 2–figure supplement 5. D). This observation supports the fact that younger paralogs derived from HMs are more likely to form HETs than older ones.

Amino acid sequence conservation could also have a direct effect on the retention of HETs, independently of the age of the duplication. For instance, among WGDs (either within true ohnologs or homeologs), which all have the same age in their own category, HM&HET pairs have higher sequence identity than HM pairs (Figure 2–figure supplement 5. B, C and E). This is also apparent for pairs of paralogs whose HM or HET structures have been solved by crystallography (Table S3). Indeed, we found that pairwise amino acid sequence identity was higher for HM&HET than for HM pairs for both entire proteins and for their binding interfaces (Figure 2. G). Furthermore, the conservation ratio of the binding interface to the non-interface regions within the available structures is higher for those forming HM&HET, suggesting a causal link between sequence identity at the interface and assembly of HM&HETs (Figure 2. H). We extended these analyses to a dataset of human paralogs (Lan and Pritchard, 2016; Singh et al., 2015) to evaluate if these trends are generalized. Whereas interfaces within PDB structures (n=65 interfaces) are more conserved than the full sequence for both HM and HM&HET motifs (Figure 2–figure supplement 6. A), we did not observe differences in the ratio of conservation of interfaces to non-interfaces (Figure 2–figure supplement 6. B). The reasons for this difference between yeast and humans remain to be explored but it could be caused by mechanisms that do not depend on interfaces to separate paralogous proteins in humans, for instance tissue-specific expression.

Considering that stable interactions are often mediated by protein domains, we looked at the domain composition of paralogs using the Protein Families Database (Pfam) (El-Gebali et al., 2019). We tested if differences in domain composition could explain the frequency of different interaction motifs. We found that 367 of 448 pairs of paralogs (82%) shared all their domain annotations (Table S3). Additionally, HM&HET paralogs tend to have more domains in common but the differences are non-significant and appear to be caused by overall sequence divergence (Figure 3–figure supplement 1. A-B). Domain gains and losses are therefore unlikely to contribute to the loss of HET complexes following the duplication of homomers.

**Figure 3:**
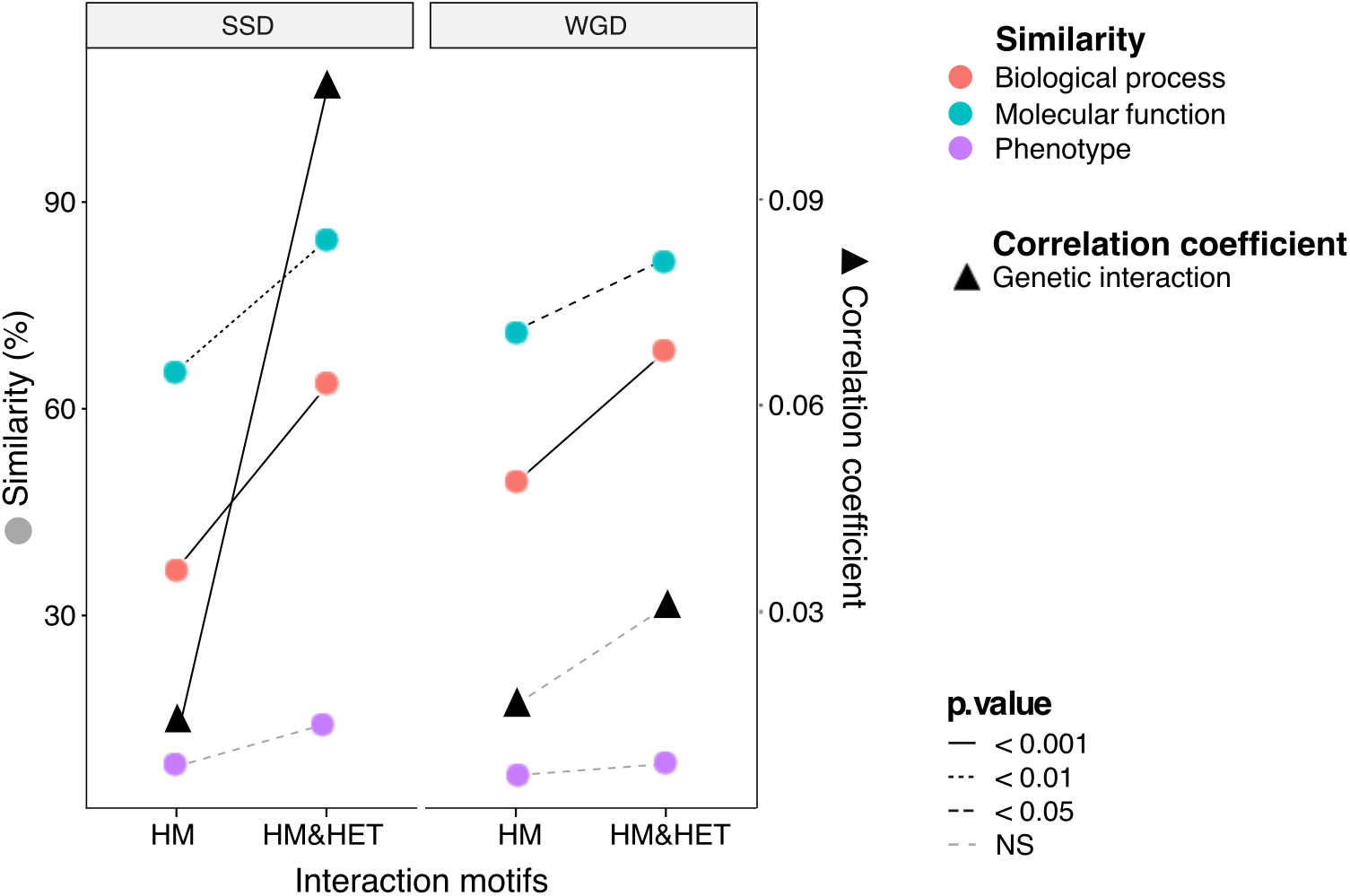
Maintenance of heteromerization between paralogs leads to greater functional similarity. The similarity score is the average proportion of shared terms (Jaccard’s index x 100) across pairs of paralogs for GO molecular functions, GO biological processes and gene deletion phenotypes. The mean values of similarity scores and of the correlation of genetic interaction profiles are compared between HM and HM&HET pairs for SSDs and WGDs. P-values are from Wilcoxon tests. Figure 3 - figure supplements 1 to 4.

### Heteromer formation correlates with functional conservation

To test if the retention of HETs correlates with the functional similarity of HM and HM&HET paralogs, we used the similarity of Gene Ontology (GO) terms, known growth phenotypes of loss-of-function mutants and patterns of genome-wide genetic interactions. These features represent the relationship of genes with cell growth and the gene-gene relationships underlying cell growth. The use of GO terms could bias the analysis because they are often predicted based on sequence features. However, phenotypes and genetic interactions are derived from unbiased experiments because interactions are tested without *a priori* consideration of a paralogs’ functions (Costanzo et al., 2016). We found that HM&HET pairs are more similar than HM for SSDs (Figure 3 and Figure 3–figure supplement 2). We observed the same trends for WGDs, although some of the comparisons are either marginally significant or non-significant (Figure 3, comparison between true ohnologs and homeologs in Figure 3–figure supplement 3). The higher functional similarity observed for HM&HET pairs could be the result of the higher sequence identity described above. However, for a similar level of sequence identity, HM&HET pairs have higher correlation of genetic interaction profiles, higher GO molecular function (for SSDs) and higher GO biological process similarity (for both SSDs and WGDs) than HM pairs (Figure 3–figure supplement 4 and GLM test in Table S7. B). Overall, the retention of HETs after the duplication of HMs appears to correlate with functional similarity, independently of sequence conservation.

### Pleiotropy contributes to the maintenance of heteromers

Since molecular interactions between paralogs predate their functional divergence, it is likely that physical association by itself affects the retention of functional similarity among paralogs. Any feature of paralogs that contributes to the maintenance of the HET state could therefore have a strong impact on the fate of new genes emerging from the duplication of HMs. A large fraction of HMs and HETs use the same binding interface (Bergendahl and Marsh, 2017), so mutations at the interface may have pleiotropic effects on both HMs and HETs (Figure 1) and correlated responses to selection. If we assume that HMs need to self-interact in order to perform their function, it is expected that natural selection would favor the maintenance of self-assembly. Negative selection on HM interfaces would act on their pleiotropic residues and thus also preserve HET interfaces, preventing the loss of HETs as a correlated response.

We tested this correlated selection model using *in silico* evolution of HM and HET protein complexes (Figure 4. A). We used a set of six representative high-quality structures of HMs (Dey et al., 2018). We evolved these HM complexes by duplicating them and following the binding energies of the resulting two HMs and HET. We let mutations occur at the binding interface 1) in the absence of selection (neutral model) and 2) in the presence of negative selection maintaining only one HM or 3) both HMs. In these three cases, we applied no selection on binding energy of the HET. In the fourth scenario, we apply selection on the HET but not on the HMs to examine if selection maintaining the HET could also favor the maintenance of HMs. Mutations that have deleterious effects on the complex under selection were lost or allowed to fix with exponentially decaying probability depending on the fitness effect (see methods) (Figure 4. A).

**Figure 4:**
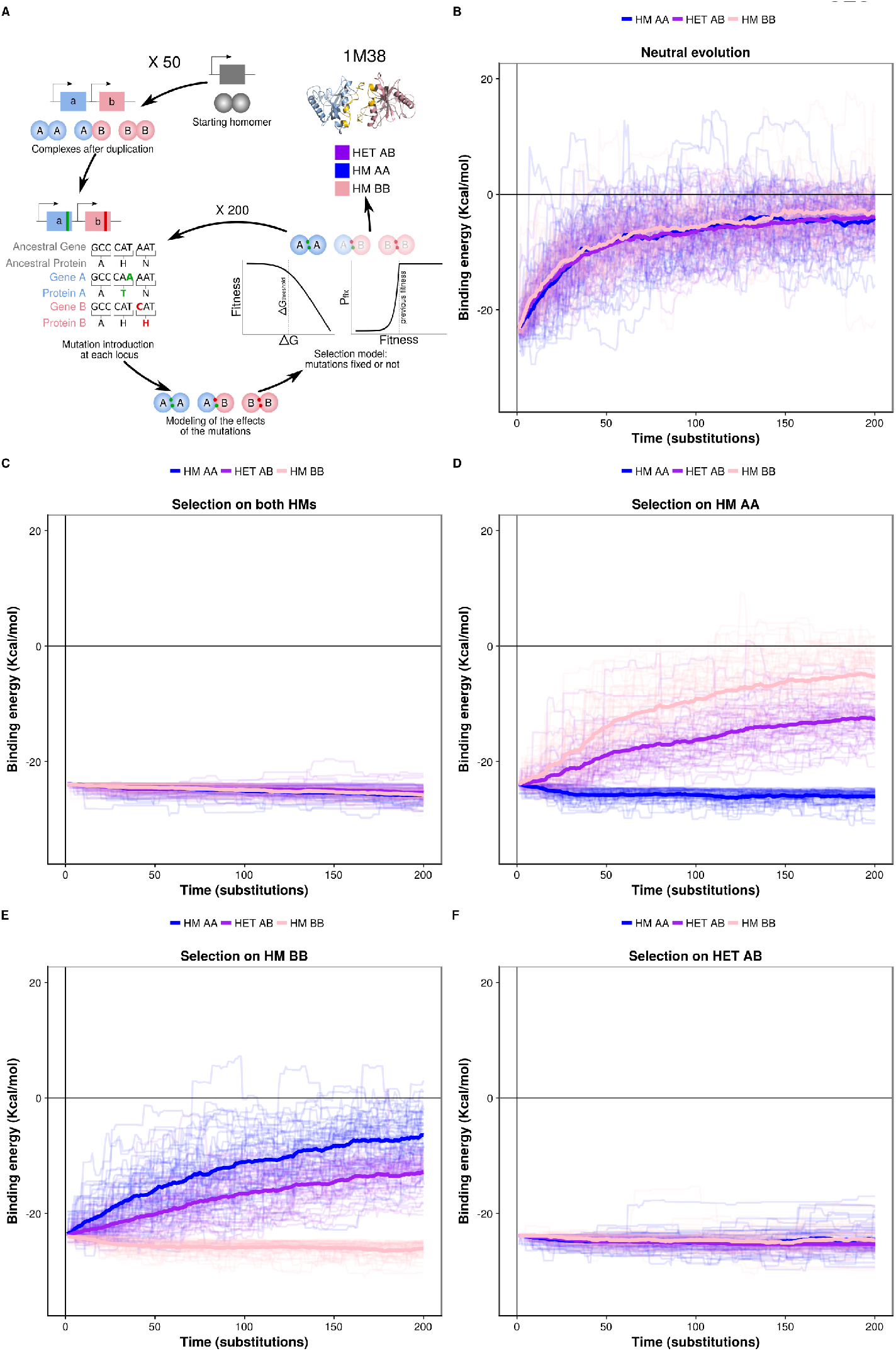
Negative selection to maintain homomers also maintains heteromers. (**A**) The duplication of a gene encoding a homomeric protein and the evolution of the complexes is simulated by applying mutations to the corresponding subunits A and B. Only mutations that would require a single nucleotide change are allowed and stop codons are disallowed. After introducing mutations, the selection model is applied to complexes and mutations are fixed or lost. (**B to F**) The binding energy of the HMs and the HET resulting from the duplication of a HM (PDB: 1M38) is followed through time under different selection regimes applied on protein stability and binding energy. More positive values indicate less favorable binding and more negative values indicate more favorable binding. (**B**) Accumulation and neutral fixation of mutations. (**C**) Selection on both HMs while the HET evolves neutrally. (**D**) Selection on HET while the HMs evolve neutrally. (**E**) Selection on HM AA or (**F**) HM BB: selection maintains one HM while the HET and the other HM evolve neutrally. Mean binding energies among replicates are shown in thick lines and the individual replicates are shown with thin lines. Fifty replicate populations are monitored in each case and followed for 200 substitutions. PDB structure 1M38 was visualized with PyMOL (Schrödinger, 2015). The number of substitutions that are fixed on average during the simulations are shown in Table S8. Figure 4 - figure supplements 1 to 4.

We find that neutral evolution leads to the destabilization of all complexes derived from the simulated duplication of a HM (PDB: 1M38) (Figure 4. B), as is expected given that there are more destabilizing mutations than stabilizing ones (Brender and Zhang, 2015; Guerois et al., 2002). Selection to maintain one HM or both HMs significantly slows down the loss of the HET with respect to the neutral scenario (Figure 4. C-E). Interestingly, the HET is being destabilized more slowly than the second HM when only one HM is under negative selection. The difficulty of losing the HET in the simulations could explain why for some paralog pairs, only one HM and the HET are preserved, as well as why there are few pairs of paralogs that specifically lose the HET (Figure 4–figure supplement 1). The reciprocal situation is also true, i.e. negative selection on HET significantly decelerates the loss of stability of both HMs (Figure 4. F). These observations hold when simulating the evolution of duplication of five other structures (Figure 4–figure supplement 2) and when simulating evolution under different combinations of parameters controlling the efficiency of selection and the length of the simulations (Figure 4–figure supplement 3). By examining the effects that single mutants (only one of the loci gets a nonsynonymous mutation) have on HMs and HET, we find that, as expected, their effects are strongly correlated and thus highly pleiotropic (Pearson’s r between 0.64 and 0.9 (Figure 4-figure supplement 4)). We observe strong pleiotropic effects of mutations for the six structures tested, which can explain the correlated responses to selection in the *in silico* evolution. Additionally, mutations tend to have greater effects on the HM than on the HET (Figure 4-figure supplement 4), which agrees with observations on HMs having a greater variance of binding energies than HETs (André et al., 2008; Lukatsky et al., 2007, 2006). As a consequence, HMs that are not under selection in our simulations show higher variability in their binding energy than HETs that are not under selection.

We examined the effects of double mutants (the two loci get a non-synonymous mutation at the interface) on HET formation to study how epistasis may influence the maintenance or loss of HET and HMs when the former or the latter are under selection. We defined epistatic effects as deviations between the observed and the expected effects of mutations on binding energy. Expected effects on HETs were calculated as the average of the effects on the HMs, which have each two subunits with the same mutation (Figure 5-figure supplement 2). We defined positive epistasis as cases where observed binding is stronger than expected (more negative ΔΔG) and negative when the effect is a reduced binding (more positive ΔΔG) compared to the expectation. In terms of evolutionary responses, positive epistasis would contribute to the retention of the HET because mutations that are slightly destabilizing HMs and thus tolerated under selection for HM stability would have less destabilizing effects on HET, slowing down its loss. On the other hand, negative epistasis could lead to the faster loss of HMs when the HET is under selection because slightly destabilizing and tolerated mutations on the HET would have stronger effects on the HMs.

**Figure 5:**
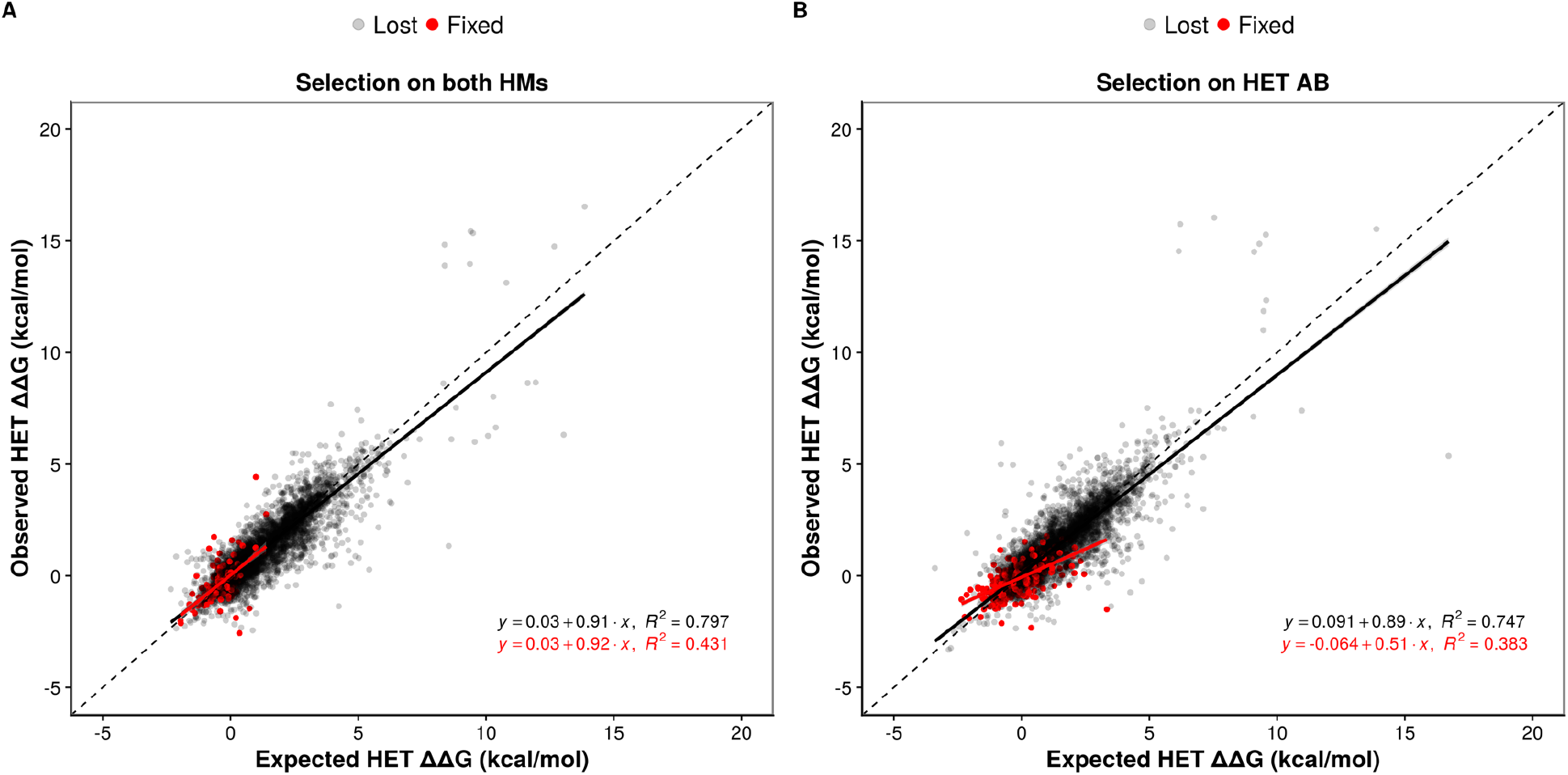
Epistasis favors the maintenance of HETs and the loss of HMs. (**A and B**) Observed effects of double mutants on HET (y-axis) are compared to their expected effects (x-axis) based on the average of their effects on the HMs when selection is applied on both HMs (n = 6777 pairs of mutations) (**A**) or on the HET (n = 6760 pairs of mutations) (**B**). Dashed lines indicate the diagonal for perfect agreement between observations and expectations (no epistasis), black regression lines indicate the best fit for the lost mutants, and red regression lines indicate the best fit for the fixed mutants. Data were obtained from simulations with PDB structure 1M38. The regression coefficients, intercepts and R2 values are indicated on the figure for fixed and lost mutations. A regression coefficient lower than one means that pairs of mutations have a less destabilizing effects on the HET than expected based on their average effects on the HMs. Figure 5 - figure supplements 1 to 3.

Regardless of the selection scenario, the mutations sampled are slightly enriched for positive epistasis, since the slopes of regression models are smaller than one (0.91 and 0.89 under selection on HMs and HET respectively). When the HMs are maintained by selection, this slightly positive epistasis is also visible in the mutations that are fixed because the epistatic effects are not selected upon. This results in a similar slope for the selected mutations as for rejected ones. Positive epistasis may therefore contribute to the maintenance of the HET (Figure 5. A). On the other hand, selection on the HET results in a further enrichment of mutations with positive epistasis (slope = 0.51, Figure 5. B). In this case, mutations tolerated in the HETs and thus fixed are more destabilizing to the HMs. This is also visible in the higher number of fixed substitutions (Table S8) when selection acts on the HET than when it acts on both HMs, particularly for mutations having opposite effects on the HMs (Figure 5–figure supplement 3). This is also manifested in significantly stronger positive epistasis among fixed pairs of mutations when the HET is under negative selection (t-test, p-value = 0.009). These observations suggest that epistasis may make HETs more robust to mutations than HMs with respect to protein complex assembly, contributing to their maintenance when the HMs are under selection and contributing to the loss of HMs when HET is under negative selection. This effect is visible in our simulations since selection on the HET results in a slow destabilization of the two HMs (Figure 4, Figure 4-figure supplement 2), especially when more mutations are attempted (Figure 4-figure supplement 3).

### Regulatory evolution may break down molecular pleiotropy

The results from simulations show that the loss of HET after the duplication of a HM occurs at a slow rate if HMs are maintained by selection and that specific rare mutations may be required for HETs to be destabilized. However, the simulations only consider the evolution of binding interfaces, which limits the modification of interactions to a subset of all mutations that can ultimately affect PPIs (Hochberg et al., 2018). Other mechanisms could involve transcriptional regulation or cell compartment localization such that paralogs are not present at the same time or in the same cell compartment. To test how regulatory evolution affects interactions, we measured the correlation coefficient of expression profiles of paralogs using mRNA microarray measurements across more than 1000 growth conditions (Ihmels et al., 2004). These expression profiles are more correlated for both SSD and WGD paralogs forming HM&HET than for those forming only HM (p-value = 6.5e-03 and 6.1e-03 respectively, Figure 6. A). This result holds using available single-cell RNAseq data (Gasch et al., 2017) although the trend is not significant for WGDs (Figure 6–figure supplement 1 A). Because we found that sequence identity was correlated with both the probability of observing HM&HET and the co-expression of paralogs, we tested if co-expression had an effect on HET formation when controlling for sequence identity. For SSDs, co-expression shows significant effects on HM&HET formation (Figure 6. C, Figure 6–figure supplement 1 B. and Table S7. B) but not for WGDs (Figure 6. C, Figure 6–figure supplement 1 B. and Table S7. B). This is true also when considering the two origins of WGDs separately (Figure 6-figure supplement 3. A-F). The differences of expression correlation between HM and HM&HET could be caused by *cis* regulatory divergence, for instance, HM&HET pairs might have more similar transcription factor binding sites. While we do observe a marginally higher transcription factor binding site similarity for HM&HET pairs than for HM pairs, the tendency is not significant, suggesting other causes for the divergence and similarity of expression profiles (Figure 6. B, Figure 6–figure supplement 2 and Table S7. B).

**Figure 6:**
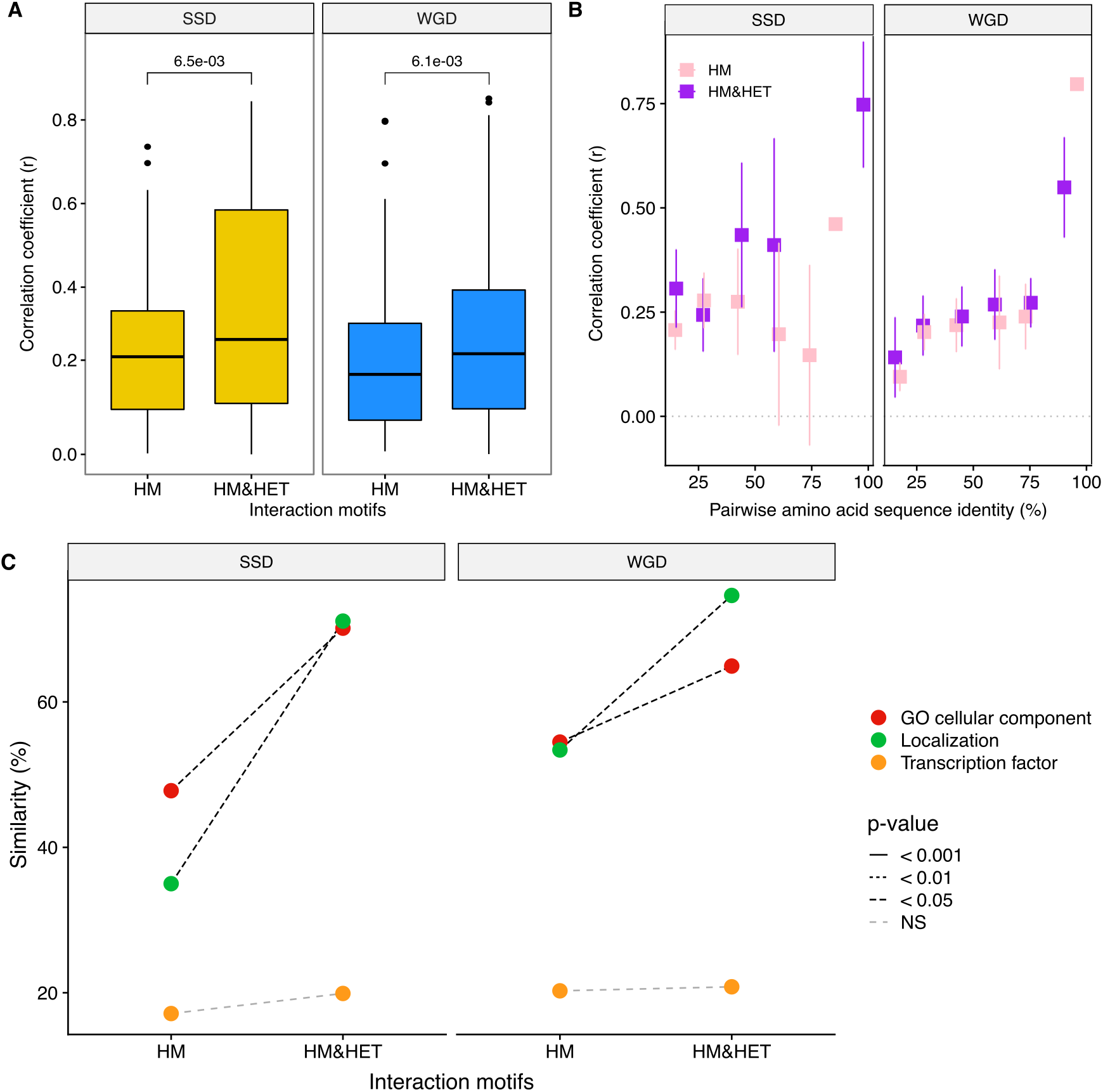
Loss of heteromerization between paralogs may result from regulatory divergence. (**A**) Correlation coefficients (Spearman’s r) between the expression profiles of paralogs. The data derives from mRNA relative expression across 1000 growth conditions (Ihmels et al., 2004). HM and HM&HET are compared for SSDs (yellow) and WGDs (blue). P-values are from t-tests. (**B**) Correlation of expression profiles between paralogs forming only HM (pink) or HM&HET (purple) as a function of their amino acid sequence identity. The data was binned into six equal categories for representation only. (**C**) Similarity of GO cellular component, GFP-based localization, and transcription factor binding sites (100* Jaccard’s index) are compared between HM and HM&HET for SSDs and WGDs. P-values are from Wilcoxon tests. Figure 6 - figure supplements 1 to 4.

Finally, we find that HM&HET paralogs are more similar than HM for both SSDs and WGDs in terms of cellular compartments (GO) and cellular localization derived from experimental data (Figure 6. C, Figure 6–figure supplement 2. B and C). For a similar level of sequence identity, HM&HET pairs have higher cellular compartment and cellular localization similarity (for both SSDs and WGDs) than HM pairs (Figure 6–figure supplement 4 and GLM test in Table S7. B). The same tendencies are observed when considering the two classes of WGDs separately (Figure 6–figure supplement 3. G-I).

Overall, coexpression, localization and GO cellular component comparison results suggest that changes in gene and protein regulation could prevent the interaction between paralogs that derive from ancestral HMs, reducing the role of structural pleiotropy in maintaining their associations.

## Discussion

Upon duplication, the properties of proteins are inherited from their ancestors, which may affect how paralogs subsequently evolve. Here, we examined the extent to which physical interactions between paralogs are preserved after the duplication of HMs and how these interactions affect functional divergence. Using reported PPI data, crystal structures and new experimental data, we found that paralogs originating from ancestral HMs are more likely to functionally diverge if they lost their ability to form HETs. We propose that non-adaptive mechanisms could play a role in the retention of physical interactions and in turn, impact on functional divergence. By developing a model of *in silico* evolution of PPIs, we found that molecular pleiotropic and epistatic effects of mutations on binding interfaces can constrain the maintenance of HET complexes even if they are not under selection. We hypothesize that this non-adaptive constraint could play a role in slowing down the divergence of paralogs but that it could be counteracted at least partly by regulatory evolution.

The proportions of HMs and HETs among yeast paralogs were first studied more than 15 years ago (Wagner, 2003). It was then suggested that most paralogs forming HETs do not have the ability to form HMs and thus, that evolution of new interactions was rapid. Since then, many PPI experiments have been performed (Chatr-Aryamontri et al., 2017; Kim et al., 2019; Stark et al., 2006; Stynen et al., 2018) and the resulting global picture is different. We found that most of the paralogs forming HETs also form HMs, suggesting that interactions between paralogs are inherited rather than gained *de novo*. This idea is supported by models predicting interaction losses to be much more likely than interaction gains after gene duplication (Gibson and Goldberg, 2009; Presser et al., 2008). Accordingly, the HM&HET state can be more readily achieved by the duplication of an ancestral HM than by the duplication of a monomeric protein followed by the gain of the HMs and of the HET. Interacting paralogs are therefore more likely to derive from ancestral HMs, as also shown by (Diss et al., 2017) using limited comparative data. For some pairs of *S. cerevisiae* paralogs presenting the HM&HET motif, we indeed detected HM formation of their orthologs from pre-whole-genome duplication species, supporting the model by which self-interaction and cross-interactions are inherited from the duplication. We did not detect HMs for both pre-whole-genome duplication species, which may reflect the incorrect expression of these proteins in *S. cerevisiae* rather than their lack of interaction.

We observed an enrichment of HMs among yeast duplicated proteins compared to singletons, as reported in previous studies (Ispolatov et al., 2005; Pereira-Leal et al., 2007; Pérez-Bercoff et al., 2010; Yang et al., 2003). Also, analyses of PPIs from large-scale experiments have shown that interactions between paralogous proteins are more common than expected by chance (Ispolatov et al., 2005; Musso et al., 2007; Pereira-Leal et al., 2007). Several adaptive hypotheses have been suggested to explain the over-representation of interacting paralogous proteins. For instance, HMs may be preferentially retained, over other duplicates, due to their ability as a source of new adaptive traits by gaining novel functions (neofunctionalization) or by splitting the original ones (subfunctionalization). For example, symmetrical HM proteins could have key advantages over monomeric ones for protein stability and regulation (André et al., 2008; Bergendahl and Marsh, 2017). Levy and Teichmann (Levy and Teichmann, 2013) suggested that the duplication of HM proteins serves as a seed for the growth of protein complexes. These duplications would allow the diversification of complexes by the asymmetric gain or loss of interactions, which would ultimately lead to the specialization of the duplicates. It is also possible that the presence of HETs itself offers a rapid way to evolve new functions. Examples include bacterial multidrug efflux transporters (Boncoeur et al., 2012) and regulatory mechanisms that evolved this way (Baker et al., 2013; Bridgham et al., 2008; De Smet et al., 2013; Kaltenegger and Ober, 2015). Finally, Natan et al. (Natan et al., 2018) showed that cotranslational folding can be a problem for homomeric proteins because of premature assembly, particularly for proteins with interfaces closer to their N-terminus. The replacement of such HMs by HETs could solve this issue by separating the translation of the proteins to be assembled on two distinct mRNAs.

Non-adaptive mechanisms could also be at play to maintain HETs. Our simulated evolution of the duplication of HMs leads to the proposal of a simple mechanism for the maintenance of HET that does not require adaptive mechanisms. A large fraction of HMs and HETs use the same binding interface (Bergendahl and Marsh, 2017) and as a consequence, negative selection on HM interfaces will also preserve HET interfaces. Our results show that mutations have correlated effects on HM and HET, which slows down the divergence of these complexes. Diss et al. (Diss et al., 2017) also suggested a non-adaptive mechanism for the maintenance of HET by showing that in the absence of their paralog, some proteins are unstable and lose their capacity to interact with other proteins. Notably, these proteins are enriched for paralogs forming HET, suggesting that the individual proteins depend on each other through these physical interactions (Diss et al., 2017). Independent observations by (DeLuna et al., 2010) also showed that the deletion of a paralog was sometimes associated with the degradation of the sister copy, particularly among HET paralogs. The Diss et al. and DeLuna et al. observations led to the proposal that paralogs could accumulate complementary degenerative mutations at the structural level after the duplication of a HM (Diss et al., 2017; Kaltenegger and Ober, 2015). This scenario would lead to the maintenance of the HET because destabilizing mutations in one subunit can be compensated by stabilizing mutations in the other, keeping binding energy and overall stability near the optimum. While compensatory mutations could also occur at different positions within identical subunits of the HMs (Uguzzoni et al., 2017), the HET would have access to those same mutations plus combinations of mutations in the two paralogous genes. As a result, the number of available compensatory mutations for the HET would be higher than that for the HM.

Furthermore, FoldX in our simulations predicts a slight overall enrichment towards positive epistasis for mutations affecting the two genes whose effects are combined in the HET. This would also contribute to the retention of the HET without adaptive mutations. Together, the smaller effect sizes of individual mutations on HET, the expanded number of compensatory mutations, and the mutational bias toward positive epistasis for the HET observed in our simulations suggest that the assembly of HET might be more robust to mutations than that of HMs. Thus, our simulations show higher potential for the specific retention of the HET than for the specific retention of the two HMs. The next step will be to text these models experimentally.

One of our observations is that WGDs present proportionally more HM&HET motifs than SSDs. We propose that this is at least partly due to the age of paralogs, which would lead to more divergence. This proposal was based on the fact that SSDs in yeast show lower sequence conservation and are thus likely older than WGDs and that even among WGDs, homeologs show less frequent HM&HET than HMs compared to true ohnologs, which are by definition younger. However, the mode of duplication itself could also impact HET maintenance. For instance, upon a whole-genome duplication event, all subunits of complexes are duplicated at the same time, which may contribute to the increased retention of WGDs in complexes compared to SSDs and thus maintain HETs. Indeed, small-scale duplications perturb the stoichiometry of complexes whereas whole-genome duplication preserves it (Birchler and Veitia, 2012; Hakes et al., 2007; Papp et al., 2003; Rice and McLysaght, 2017). In addition, Fares et al. (Fares et al., 2013) suggested that SSDs display higher evolutionary rates than WGDs, which could lead to the loss of their interactions. Another factor that differs is that some WGDs are maintained due to selection for higher gene dosage (Ascencio et al., 2017; Edger and Pires, 2009; Gout and Lynch, 2015; Sugino and Innan, 2006; Thompson et al., 2016). Therefore, the ancestral gene sequence, regulation and function are conserved, which ultimately favors the maintenance of HETs among WGDs.

We noticed a significant fraction of paralogs forming only HMs but not HET, including some cases of recent duplicates, indicating that the forces maintaining HETs can be overcome. Moreover, although SSDs are more divergent than WGDs on average, the sequence divergence and domain composition differ slightly (not significant) between HMs and HM&HETs, suggesting a mechanism other that amino acid sequence divergence for HET loss. Duplicate genes in yeast and other model systems often diverge quickly in terms of transcriptional regulation (Li et al., 2005; Thompson et al., 2013) due to *cis* regulatory mutations (Dong et al., 2011). Because transcriptional divergence of paralogs can directly change PPI profiles, expression changes would be able to rapidly change a motif from HM&HET to HM. Indeed, Gagnon-Arsenault et al. (Gagnon-Arsenault et al., 2013) showed that switching the coding sequences between paralogous loci was sometimes sufficient to change PPI specificity in living cells. Protein localization can also be an important factor affecting the ability of proteins to interact (Rochette et al., 2014). We found that paralogs that derive from HMs and that have lost their ability to form HETs are less co-regulated and less co-localized. This divergence suggests that regulatory evolution could play a role in relieving duplicated homomeric proteins from the correlated effects of mutations affecting shared protein interfaces.

Overall, our analyses show that the duplication of self-interacting proteins creates paralogs whose evolution is constrained by pleiotropy in ways that are not expected for monomeric paralogs. Pleiotropy has been known to influence the architecture of complex traits and thus to shape their evolution (Wagner and Zhang, 2011). However, how it takes place at the molecular level and how it can be overcome to allow molecular traits to evolve independently is still largely unknown. Here, we provide a simple system in which the role of pleiotropy can be examined at the molecular level. Because gene duplication is a major mechanism responsible for the evolution of cellular networks and because a large fraction of proteins are oligomeric, the pleiotropic and epistatic constraints described here could be an important force in shaping protein networks. Another important result is that negative selection for the maintenance of heteromers of paralogs is not needed for their preservation on the long term, further enhancing the role of non-adaptive evolution in shaping the complexity of cellular structures (Lynch et al., 2014).

## Material and Methods

The protein-protein interactions identified in this publication have been submitted to the IMEx (http://www.imexconsortium.org) consortium through IntAct (Orchard et al., 2014) and are assigned the identifier IM-26944. All scripts used to analyze the data are available at https://github.com/landrylaboratory/Gene_duplication_2019.

### 1. Characterization of paralogs in *S. cerevisiae* genome

#### 1.1 Classification of paralogs by mechanism of duplication

We classified duplicated genes in three categories according to their mechanism of duplication: small-scale duplicates (SSDs); whole-genome duplicates (WGDs) (Byrne and Wolfe, 2005); and double duplicates (2D, SSDs and WGDs). We removed WGDs from the paralogs defined in (Guan et al., 2007) to generate the list of SSDs. Among paralog pairs with less than 20% of sequence identity in the multiple sequence alignments (data from MSA, (Edgar, 2004)), we kept only those sharing the same phylome (PhylomeDB (Huerta-Cepas et al., 2008)) to make sure they were true paralogs. If one of the two paralogs of an SSD pair is associated to another paralog in a WGD pair, this paralog was considered a 2D (Tables S1 and S2). To decrease the potential bias from multiple duplication events, we removed the 2Ds and paralogs from successive small-scale genome duplications from the data on interaction motifs. We used data from (Marcet-Houben and Gabaldón, 2015) to identify WGDs that are likely true ohnologs or that originated from allopolyploidization (homeologs).

#### 1.2 Sequence similarity

Conversion tables between PhylomeDB IDs and systematic yeast IDs were downloaded from ftp://phylomedb.org/phylomedb/all_id_conversion.txt.gz on May 15th, 2019. Sequence identity was calculated from multiple sequence alignments from phylome 0003 from PhylomeDB (Huerta-Cepas et al., 2008). The yeast phylome consists of 60 completely sequenced fungal species, with *Homo sapiens* and *Arabidopsis thaliana* as outgroups. Sequences in these phylomes were aligned with MUSCLE v 3.6. When two paralogs were not found in the same multiple sequence alignment from PhylomeDB (32 pairs out of 462 pairs), the sequences were taken from the reference proteome of *S. cerevisiae* assembly R64-1-1 downloaded on April 16th, 2018 from the Ensembl database at (http://useast.ensembl.org/info/data/ftp/index.html) (Zerbino et al., 2018) and realigned to the rest of the phylome with MUSCLE version 3.8.31 (Edgar, 2004). For six pairs of paralogs that did not have phylomeDB IDs assigned to them, pairwise alignments of their sequences with MUSCLE version 3.8.31 (Edgar, 2004) were used.

#### 1.3 Function, transcription factor binding sites, localization protein complexes, and Pfam annotations

We obtained GO terms (GO slim) from SGD (Cherry et al., 2012) in September 2018. We removed terms corresponding to missing data and created a list of annotations for each SSD and WGD gene. Annotations were compared to measure the extent of similarity between two members of a pair of duplicates. We calculated the similarity of molecular function, cellular component and biological process taking the number of GO terms in common divided by the total number of unique GO terms of the two paralogs combined (Jaccard index). We compared the same way transcription factor binding sites using YEASTRACT data (Teixeira et al., 2018, 2006), cellular localizations extracted from YeastGFP database (Huh et al., 2003) and many phenotypes associated with the deletion of paralogs (data from SGD in September 2018). For the deletion phenotypes, we kept only information with specific changes (a feature observed and a direction of change relative to wild type). We compared the pairwise correlation of genetic interaction profiles using the genetic interaction profile similarity (measured by Pearson’s correlation coefficient) of non-essential genes available in TheCellMap database (version of March 2016) (Usaj et al., 2017). We used the median of correlation coefficients if more than one value was available for a given pair. Non-redundant set of protein complexes was derived from the Complex Portal (Meldal et al., 2015), the CYC2008 catalogue (Pu et al., 2009, 2007) and Benschop et al., (Benschop et al., 2010).

We downloaded Pfam domain annotations (El-Gebali et al., 2019) for the whole *S. cerevisiae* reference proteome on May 2nd, 2019 from the UniprotKB database (The UniProt Consortium, 2019). We removed pairs of paralogs for which at least one of the proteins had no annotated domains and calculated the Jaccard index (Table S3).

### 2. HMs and HETs identified from databases

To complement our experimental data, we extracted HMs and HETs published in BioGRID version BIOGRID-3.5.166 (Chatr-Aryamontri et al., 2017, 2013). We used data derived from the following detection methods: Affinity Capture-MS, Affinity Capture-Western, Reconstituted Complex, Two-hybrid, Biochemical Activity, Co-crystal Structure, Far Western, FRET, Protein-peptide, PCA and Affinity Capture-Luminescence.

It is possible that some HMs or HETs are absent from the database because they have been tested but not detected. This negative information is not reported in databases. We therefore attempted to discriminate non-tested interactions from truly non interacting pairs. A study in which there was not a single HM reported was considered as missing data for all HMs. For both HMs and HETs, the presence of a protein (or both proteins for HET) as both bait and prey but the absence of interaction was considered as evidence for no interaction. Otherwise, it was considered as missing data (coded NA).

We also considered data from crystal structures. If a HM was detected in the Protein Data Bank (PDB) (Berman et al., 2000), we inferred that it was present. If the HM was not detected but the monomer was reported, it is likely that there is no HM for this protein and it was thus considered non-HM. If there was no monomer and no HM, the data were considered as missing. We proceeded the same way for HETs.

Data on genome-wide HM screens was obtained from (Kim et al., 2019; Stynen et al., 2018). The two methods relied on Protein-fragment complementation assays (PCA), the first one using the dihydrofolate reductase (DHFR) enzyme as a reporter and the second one, a fluorescent protein (also known as Bimolecular fluorescence complementation (BiFC)). We discarded proteins from (Stynen et al., 2018) flagged as problematic by (Rochette et al., 2014; Stynen et al., 2018; Tarassov et al., 2008) and false positives identified by (Kim et al., 2019). All discarded data was considered as missing data. We examined all proteins tested and considered them as HM if they were reported as positive and as non-HM if tested but not reported as positive.

### 3. Experimental Protein-fragment complementation assay

We performed a screen using PCA based on DHFR (Tarassov et al., 2008) following standard procedures (Rochette et al., 2014; Tarassov et al., 2008). The composition of all following media used in this study is described in Table S11.

#### 3.1 DHFR strains

We identified 485 pairs of SSDs and 156 pairs of WGDs present in the Yeast Protein Interactome Collection (Tarassov et al., 2008) and another set of 155 strains constructed by (Diss et al., 2017). We retrieved strains from the collection (Tarassov et al., 2008) and we let them grow on NAT (DHFR F[1,2] strains) and HygB (DHFR F[3] strains) media. We confirmed the insertion of the DHFR fragments at the correct location by colony PCR using a specific forward Oligo-C targeting a few hundred base pairs upstream of the fusion and a reverse complement oligonucleotide ADHterm_R located in the ADH terminator after the DHFR fragment sequence (Table S11). Cells from colonies were lysed in 40 μL of 20 mM NaOH for 20 min at 95°C. Tubes were centrifuged for 5 min at 1792 g and 2.5 μL of supernatant was added to a PCR mix composed of 16.85 μL of DNAse free water, 2.5 μL of 10X Taq buffer (BioShop Canada Inc., Canada), 1.5 μL of 25 mM MgCl2, 0.5 μL of 10 mM dNTP (Bio Basic Inc., Canada), 0.15 μL of 5 U/μL Taq DNA polymerase (BioShop Canada Inc., Canada), 0.5 μL of 10 μM Oligo-C and 0.5 μL of 10 μM ADHterm_R. The initial denaturation was performed for 5 min at 95°C and was followed by 35 cycles of 30 sec of denaturation at 94°C, 30 sec of annealing at 55°C, 1 min of extension at 72°C and by a 3 min final extension at 72°C. We confirmed by PCR 2025 out of the 6585 strains from the DHFR collection and 126 strains out of the 154 from (Diss et al., 2017) (Tables S9, S10, and S12).

The missing or non-validated strains were constructed *de novo* using the standard DHFR strain construction protocol (Michnick et al., 2016; Rochette et al., 2015). The DHFR fragments and associated resistance modules were amplified from plasmids pAG25-linker-F[1,2]-ADHterm (NAT resistance marker) and pAG32-linker-F[3]-ADHterm (HygB resistance marker) (Tarassov et al., 2008) using oligonucleotides defined in (Table S12). pCr mix was composed of 16.45 μL of DNAse free water, 1 μL of 10 ng/μL plasmid, 5 μL of 5X Kapa Buffer (Kapa Biosystems, Inc., A Roche Company, Canada), 0.75 μL of 10 mM dNTPs, 0.3 μL of 1 U/μL Kapa HiFi HotStart DNA polymerase (Kapa Biosystems, Inc., A Roche Company, Canada) and 0.75 μL of both forward and reverse 10 μM oligos. The initial denaturation was performed for 5 min at 95°C and was followed by 32 cycles of 20 sec of denaturation at 98°C, 15 sec of annealing at 64.4°C, 2.5 min of extension at 72°C and 5 min of a final extension at 72°C.

We performed strain construction in BY4741 (MATa *his3Δ leu2Δ met15Δ ura3Δ*) and BY4742 (MATα *his3Δ leu2Δ lys2Δ ura3Δ*) competent cells prepared as in (Gagnon-Arsenault et al., 2013) for the DHFR F[1,2] and DHFR F[3] fusions, respectively. Competent cells (20 μL) were combined with 8 μL of PCR product (~0.5-1 μg/μL) and 100 μL of Plate Mixture (PEG3350 40%, 100 mM of LiOAc, 10 mM of Tris-Cl pH 7.5 and 1 mM of EDTA). Cells were vortexed and incubated at room temperature without agitation for 30 min. After adding 15 μL of DMSO and mixing thoroughly, heat shock was then performed by incubating in a water bath at 42°C for 15-20 min. Following the heat shock, cells were spun down at 400 g for 3 min. Supernatant was removed by aspiration and cell pellets were resuspended in 100 μL of YPD. Cells were allowed to recover from heat shock for 4 hours at 30°C before being plated on NAT (DHFR F[1,2] strains) or HygB (DHFR F[3] strains) plates. Cells were incubated at 30°C for 3 days. The correct integration of DHFR fragments was confirmed by colony PCR as described above and later by sequencing (Plateforme de séquençage et de génotypage des génomes, CRCHUL, Canada) for specific cases where the interaction patterns suggested a construction problem, for instance when the HET was observed in one direction only or when one HM was missing for a given pair. At the end, we reconstructed and validated 146 new strains (Tables S9 and S10). From all available strains, we selected pairs of paralogs for which we had both proteins tagged with both DHFR fragments (four different strains per pair). This resulted in 1172 strains corresponding to 293 pairs of paralogs (Tables S9 and S10). We finally discarded pairs considered as forming false positives by (Tarassov et al., 2008), which resulted in 235 pairs.

#### 3.2 Construction of DHFR plasmids for orthologous gene expression

For the plasmid-based PCA, Gateway cloning-compatible destination plasmids pDEST-DHFR F[1,2] (TRP1 and LEU2) and pDEST-DHFR F[3] (TRP1 and LEU2) were constructed based on the CEN/ARS low-copy yeast two-hybrid (Y2H) destination plasmids pDEST-AD (TRP1) and pDEST-DB (LEU2) (Rual et al., 2005). A DNA fragment having I-CeuI restriction site was amplified using DEY001 and DEY002 primers (Table S12) without template and another fragment having PI-PspI/I-SceI restriction site was amplified using DEY003 and DEY004 primers (Table S11) without template. pDEST-AD and pDEST-DB plasmids were each digested by PacI and SacI and mixed with the I-CeuI fragment (destined to the PacI locus) and PI-PspI/I-SceI fragment (destined to the SacI locus) for Gibson DNA assembly (Gibson et al., 2009) to generate pDN0501 (TRP1) and pDN0502 (LEU2). Four DNA fragments were then prepared to construct the pDEST-DHFR F[1,2] vectors: (i) a fragment containing ADH1 promoter; (ii) a fragment containing Gateway destination site; (iii) a DHFR F[1,2] fragment; and (iv) a backbone plasmid fragment. The ADH1 promoter fragment was amplified from pDN0501 using DEY005 and DEY006 primers (Table S12) and the Gateway destination site fragment was amplified from pDN0501 using DEY007 and DEY008 primers (Table S12). The DHFR-F[1,2] fragment was amplified from pAG25-linker-F[1,2]-ADHterm (Tarassov et al., 2008) using DEY009 and DEY010 primers (Table S12).

The backbone fragment was prepared by restriction digestion of pDN0501 or pDN0502 using I-CeuI and PI-PspI and purified by size-selection. The four fragments were assembled by Gibson DNA assembly where each fragment pair was overlapping with more than 30 bp, producing pHMA1001 (TRP1) or pHMA1003 (LEU2). The PstI-SacI region of the plasmids was finally replaced with a DNA fragment containing an amino acid flexible polypeptide linker (GGGGS) prepared by PstI/SacI double digestion of a synthetic DNA fragment DEY011 to produce pDEST-DHFR F[1,2] (TRP1) and pDEST-DHFR F[1,2] (LEU2). The DHFR F[3] fragment was then amplified from pAG32-linker-F[3]-ADHterm with DEY012 and DEY013 primers (Table S11), digested by SpeI and PI-PspI, and used to replace the SpeI–PI-PspI region of the pDEST-DHFR F[1,2] plasmids, producing pDEST-DHFR F[3] (TRP1) and pDEST-DHFR F[3] (LEU2) plasmids. In this study, we used pDEST-DHFR F[1,2] (TRP1) and pDEST-DHFR F[3] (LEU2) for the plasmid-based DHFR PCA. After Gateway LR cloning of Entry Clones to these destination plasmids, the expression plasmids encode protein fused to the DHFR fragments via an NPAFLYKVVGGGSTS linker.

We obtained the orthologous gene sequences for the mitochondrial translocon complex and the transaldolase proteins of *Lachancea kluyveri* and *Zygosaccharomyces rouxii* from the Yeast Gene Order Browser (YGOB) (Byrne and Wolfe, 2005). Each ORF was amplified using oligonucleotides listed in Table S11. We used 300 ng of purified PCR product to set a BPII recombination reaction (5 μL) into the Gateway Entry Vector pDONR201 (150 ng) according to the manufacturer’s instructions (Invitrogen, USA). BpII reaction mix was incubated overnight at 25°C. The reaction was inactivated with proteinase K. The whole reaction was used to transform MC1061 competent *E. coli* cells (Green and Rogers, 2013), followed by selection on solid 2YT medium supplemented with 50 mg/L of kanamycin (BioShop Inc., Canada) at 37°C. Positive clones were detected by PCR using an ORF specific oligonucleotide and a general pDONR201 primer (Table S12). We then extracted the positive Entry Clones using Presto™ Mini Plasmid Kit (Geneaid Biotech Ltd, Taiwan) for downstream application.

LRII reactions were performed by mixing 150 ng of the Entry Clone and 150 ng of expression plasmids (pDEST-DHFR F[1,2]-TRP1 or pDEST-DHFR F[3]-LEU2) according to manufacturer’s instructions (Invitrogen, USA). The reactions were incubated overnight at 25°C and inactivated with proteinase K. We used the whole reaction to transform MC1061 competent *E. coli* cells, followed by selection on solid 2YT medium supplemented with 100 mg/L ampicillin (BioShop Inc., Canada) at 37°C. Positive clones were confirmed by PCR using a ORF specific primer and a plasmid universal primer. The sequence-verified expression plasmids bearing the orthologous fusions with DHFR F[1,2] and DHFR F[3] fragments were used to transform the yeast strains YY3094 (MATa *leu2-3,112 trp1-901 his3-200 ura3-52 gal4Δ gal80*Δ *LYS2::P_GAL1_-HIS3 MET2::PGAL7-lacZ cyh2^R^ can1*Δ::*P_CMV_-rtTA-KanMX4*) and YY3095 (MATα *leu2-3,112 trp1-901 his3-200 ura3-52 gal4*Δ *gal80*Δ *LYS2::P_GAL1_-HIS3 MET2::PGAL7-lacZ cyh2^R^ can1Δ::T_ADH1_-P_tetO2_-Cre-T_CYC1_-KanMX4*), respectively. Selection was done on SC -trp -ade (YY3094) or on SC -leu - ade (YY3095). The strains YY3094 and YY3095 were generated from BFG-Y2H toolkit strains RY1010 and RY1030 (Yachie et al., 2016), respectively, by restoring their wild type *ADE2* genes. The *ADE2* gene was restored by homologous recombination of the wild type sequence cassette amplified from the laboratory strain BY4741 using primers DEY014 and DEY015 (Table S12). SC -ade plates were used to obtain successful transformants.

#### 3.3 DHFR PCA experiments

Three DHFR PCA experiments were performed, hereafter referred to as PCA1, PCA2 and PCA3. The configuration of strains on plates and the screenings were performed using robotically manipulated pin tools (BM5-SC1, S&P Robotics Inc., Toronto, Canada (Rochette et al., 2015)). We first organized haploid strains in 384 colony arrays containing a border of control strains using a cherry-picking 96-pin tool (Figure 2–figure supplement 7). We constructed four haploid arrays corresponding to paralog 1 and 2 (P1 and P2) and mating type: MATa P1-DHFR F[1,2]; MATa P2-DHFR F[1,2] (on NAT medium); MATα P1-DHFR F[3]; MATα P2-DHFR F[3] (on HygB medium). Border control strains known to show interaction by PCA (MATa *LSM8*-DHFR F[1-2] and MATα *CDC39*-DHFR F[3]) were incorporated respectively in all MATa DHFR F[1,2] and MATα DHFR F[3] plates in the first and last columns and rows. The strains were organized as described in Figure 2–figure supplement 7. The two haploid P1 and P2 384 plates of the same mating type were condensated into a 1536 colony array using a 384-pintool. The two 1536 arrays (one MATa DHFR F[1,2], one MATα DHFR F[3]) were crossed on YPD to systematically test P1-DHFR F[1,2] / P1-DHFR F[3], P1-DHFR F[1,2]/P2-DHFR F[3], P2-DHFR F[1,2]/P1-DHFR F[3] and P2-DHFR F[1,2]/P2-DHFR F[3] interactions in adjacent positions. We performed two rounds of diploid selection (S1 to S2) by replicating the YPD plates onto NAT+HygB and growing for 48 hours. The resulting 1536 diploid plates were replicated twice for 96 hours on DMSO -ade -lys - met control plates (for PCA1 and PCA2) and twice for 96 hours on the selective MTX -ade -lys - met medium (for all runs). Five 1536 PCA plates (PCA1-plate1, PCA1-plate2, PCA2, PCA3-plate1 and PCA3-plate2) were generated this way. We tested the interactions between 277 pairs in five to twenty replicates each (Table S3).

We also used the robotic platform to generate three bait and three prey 1536 arrays for the DHFR plasmid-based PCA, testing each pairwise interaction at least four times. We mated all MATa DHFR F[1,2] and MATα DHFR F[3] strains on YPD medium at room temperature for 24 hours. We performed two successive steps of diploid selection (SC -leu -trp -ade) followed by two steps on DMSO and MTX media (DMSO -leu -trp -ade and MTX -leu -trp -ade). We incubated the plates of diploid selection at 30°C for 48 hours. Finally, plates from both MTX steps were incubated and monitored for 96 hours at 30°C.

#### 3.4 Analysis of DHFR PCA results

##### 3.4.1 Image analysis and colony size quantification

All images were analysed the same way, including images from (Stynen et al., 2018). Images of plates were taken with a EOS Rebel T5i camera (Canon, Tokyo, Japan) every two hours during the entire course of the PCA experiments. Incubation and imaging were performed in a spImager custom platform (S&P Robotics Inc., Toronto, Canada). We considered images after two days of growth for diploid selection plates and after four days of growth for DMSO and MTX plates. Images were analysed using *gitter* (R package version 1.1.1 (Wagih and Parts, 2014)) to quantify colony sizes defining a square around the colony center and measuring the foreground pixel intensity minus the background pixel intensity.

##### 3.4.2 Data filtering

For the images from (Stynen et al., 2018), we filtered data based on the diploid selection plates. Colonies smaller than 200 pixels were considered as missing data rather than as non-interacting strains. For PCA1, PCA2 and PCA3, colonies flagged as irregular by *gitter* (as S (colony spill or edge interference) or S, C (low colony circularity) flags) or that did not grow on the last diploid selection step or on DMSO medium (smaller than quantile 25 minus the interquartile range) were considered as missing data. We considered only bait-prey pairs with at least four replicates and used the median of colony sizes as PCA signal. The data was finally filtered based on the completeness of paralogous pairs so we could test HMs and HETs systematically. Thus, we finally obtained results for 241 paralogous pairs (Tables S3 and S4). Median colony sizes were log2 transformed after adding a value of 1 to all data to obtain PCA scores. The results of (Stynen et al., 2018) and PCA1, PCA2 and PCA3 were strongly correlated (Figure 2–figure supplement 1. B). Similarly, the results correlate well with those reported by (Tarassov et al., 2008) (Figure 2–figure supplement 1. C).

##### 3.4.3 Detection of protein-protein interactions

The distribution of PCA scores was modeled per duplication type (SSD and WGD) and per interaction tested (HM or HET) as in (Diss et al., 2017) with the *normalmixEM* function (default parameters) available in the R mixtools package (Benaglia et al., 2009). The background signal on MTX was used as a null distribution to which interactions were compared. The size of colonies (PCA scores (PCA_s_)) were converted to z-scores using the mean (μ_b_) and standard deviation (sd_b_) of the background distribution (Z_s_ = (PCA_s_ - μ_b_)/sd_b_). PPI were considered detected if Z_s_ of the bait-prey pair was greater than 2.5 (Figure 2–figure supplement 8) (Chrétien et al., 2018).

We observed 24 cases in which only one of the two possible HET interactions was detected (P1-DHFR F[1,2] x P2-DHFR F[3] or P2-DHFR F[1,2] x P1-DHFR F[3]). It is typical for PCA assays to detect interactions in only one orientation or the other (See (Tarassov et al., 2008)). However, this could also be caused by one of the four strains having an abnormal fusion sequence. We verified by PCR and sequenced the fusion sequences to make sure this was not the case. The correct strains were conserved and the other ones were re-constructed and retested. No cases of unidirectional HET were observed in our final results. For all 71 pairs after reconstruction, both reciprocal interactions were detected.

##### 3.4.4 Dataset integration

The PCA data was integrated with other data obtained from databases. The overlaps among the different datasets and the results of our PCA experiments are shown in Figure 2–figure supplement 2.

### 4. Gene expression in MTX condition

#### 4.1 Cell cultures for RNAseq

We used the border control diploid strain from the DHFR PCA (MATa/α *LSM8*-DHFR F[1,2]/*LSM8 CDC39/CDC39*-DHFR F[3]) to measure expression profile in MTX condition. Three overnight pre-cultures were grown separately in 5 ml of NAT+HygB at 30°C with shaking at 250 rpm. A second set of pre-cultures were grown starting from a dilution at OD_600_ = 0.01 in 50 ml in the same condition to an OD_600_ of 0.8 to 1. Final cultures were started at OD_600_ = 0.03 in 250 ml of synthetic media supplemented with MTX or DMSO (MTX -ade -trp -leu or DMSO -ade -trp - leu) at 30°C with shaking at 250 rpm. These cultures were transferred to 5 x 50 ml tubes when they reached an OD_600_ of 0.6 to 0.7 and centrifuged at 1008 g at 4°C for 1 min. The supernatant was discarded and cell pellets were frozen in liquid nitrogen and stored at −80°C until processing. RNA extractions and library generation and amplification were performed as described in (Eberlein et al., 2019). Briefly, the Quantseq 3’ mRNA kit (Lexogen, Vienna, Austria) was used for library preparation (Moll et al., 2014) following the manufacturer’s protocol. The PCR cycles number during library amplification was adjusted to 16. The six libraries were pooled and sequenced on a single Ion Torrent chip (ThermoFisher Scientific, Waltham, United States) for a total of 7,784,644 reads on average per library. Barcodes associated to the samples in this study are listed in Table S5.

#### 4.2 RNAseq analysis

Read quality statistics were retrieved from the program FastQC (Andrews, 2010). Reads were cleaned using cutadapt (Martin, 2011). We removed the first 12 bp, trimmed the poly-A tail from the 3’ end, trimmed low-quality ends using a cutoff of 15 (phred quality + 33) and discarded reads shorter than 30 bp. The number of reads before and after cleaning can be found in Table S5. Raw sequences can be downloaded under the NCBI BioProject ID PRJNA480398.

Cleaned reads were aligned on the reference genome of S288c from SGD (S288C_reference_genome_R64-2-1_20150113.fsa version) using bwa (Li and Durbin, 2009). Because we used a 3’mRNA-Seq Library, reads mapped largely to 3’UTRs. We increased the window of annotated genes in the SGD annotation (saccharomyces_cerevisiae_R64-2-1_20150113.gff version) using the UTR annotation from (Nagalakshmi et al., 2008). Based on this reference genes-UTR annotation, the number of mapped reads per genes was estimated using htseq-count of the Python package HTSeq (Anders et al., 2015) and reported in Table S6.

#### 4.3 Correlation of gene expression profiles

The correlation of expression profiles for paralogs was calculated using Spearman’s correlation from large-scale microarray data (Ihmels et al., 2004) over 1000 mRNA expression profiles from different conditions and different cell cycle phases. These results were compared and confirmed with a large-scale expression data from normalized RNAseq single cells of *S. cerevisiae* grown in normal or stressful conditions (0.7 M NaCl) and from different cell cycle phases (Gasch et al., 2017).

### 5. Structural analyses

#### 5.1 Sequence conservation in binding interfaces of yeast complexes

##### 5.1.1 Identification of crystal structures

The sequences of paralogs classified as SSDs or WGDs (Byrne and Wolfe, 2005; Guan et al., 2007) were taken from the reference proteome of *Saccharomyces cerevisiae* assembly R64-1-1 and searched using BLASTP (version 2.6.0+) (Camacho et al., 2009) to all the protein sequences contained in the Protein Data Bank (PDB) downloaded on September 21^st^, 2017 (Berman et al., 2000). Due to the high sequence identity of some paralogs (up to 95%), their structures were assigned as protein subunits from the PDB that had a match with 100% sequence identity and an E-value lower than 1e-6. Only crystal structures that spanned more than 50% of the full protein length were kept for the following analyses. The same method was used to retrieve PDB structures for human paralogous proteins. The human reference proteome Homo_sapiens.GRCh38.pep.all.fa was downloaded on May 16th, 2019 from the Ensembl database (http://useast.ensembl.org/info/data/ftp/index.html) (Zerbino et al., 2018). Pairs of paralogs were retrieved from two different datasets (Lan and Pritchard, 2016; Singh et al., 2015). Protein interactions for those proteins were taken from a merged dataset from the BioGRID (Chatr-Aryamontri et al., 2017) and IntAct (Orchard et al., 2014) databases. The longest protein isoforms for each gene in the dataset were aligned using BLASTP to the set of sequences from the PDB. Matches with 100% sequence identity and E-values below 1e-6 were assigned to the subunits from the PDB structures.

##### 5.1.2 Identification of interfaces

Residue positions involved in protein binding interfaces were defined based on the distance of residues to the other subunit (Tsai et al., 1996). Contacting residues are defined as those whose two closest non-hydrogen atoms are separated by a distance smaller than the sum of their van der Waals radii plus 0.5 Å. Reference van der Waals radii were obtained with FreeSASA version 2.0.1 (Mitternacht, 2016). Nearby residues are those whose alpha carbons are located at a distance smaller than 6 Å. All distances were measured using the Biopython library (version 1.70) (Cock et al., 2009).

##### 5.1.3 Sequence conservation within interfaces

The dataset of PDB files was then filtered to include only the crystallographic structures with the highest resolution available for each complex involving direct contacts between subunits of paralogs. Full-length protein sequences from the reference proteome were then aligned to their matching subunits from the PDB with MUSCLE version 3.8.31 (Edgar, 2004) to assign the structural data to the residues in the full-length protein sequence. These full-length sequences were then aligned to their paralogs and sequences from PhylomeDB phylogenies (phylome 0003) (Huerta-Cepas et al., 2008) with MUSCLE version 3.8.31. Only three pairs of paralogs that needed realignment were included in this analysis. Sequence identity was calculated within interface regions, which considered the contacting and nearby residues. Paralogs were classified as HM or HM&HET based on the data shown in Table S3. PDB identifiers for structures included in this analysis are shown in Table S13. Pairs of paralogs for which the crystallized domain was only present in one of the proteins were not considered for this analysis.

A similar procedure was applied to the human proteins, with sequences aligned to their corresponding PhylomeDB phylogenies from phylome 0076 (A new human phylome release using current phylogenetic pipeline with updated proteomes) resulting from forward and reverse alignments obtained with MUSCLE 3.8, MAFFT v6.712b and DIALIGN-TX, and merged with M-COFFEE (Huerta-Cepas et al., 2008). Considering that human genes code for multiple isoforms, we took the isoforms from the two paralogs that had the highest sequence identity with respect to the PDB structure. When a gene coded for multiple isoforms that were annotated with identical protein sequence in the human reference proteome, we only kept one of them. This resulted in a set of 40 HM interfaces and 25 HM&HET interfaces for a total of 54 different pairs (35 HM pairs and 19 HM&HET). Pairs of paralogs were classified as HM or HM&HET based on the data in Tables S14 and S15.

#### 5.2 Simulations of coevolution of protein complexes

##### 5.2.1 Mutation sampling during evolution of protein binding interfaces

Simulations were carried out with high quality crystal structures of homodimeric proteins from PDB (Berman et al., 2000). Four of them (PDB: 1M38, 2JKY, 3D8X, 4FGW) were taken from the above data set of structures that matched yeast paralogs and two others from the same tier of high quality structures (PDB: 1A82, 2O1V). The simulations model the duplication of the gene encoding the homodimer, giving rise to separate copies that can accumulate different mutations, leading to the formation of HMs and HETs as in Figure 1.

Mutations were introduced using a transition matrix whose substitution probabilities consider the genetic code and allow only substitutions that would require a single base change in the underlying codons (Thorvaldsen, 2016). Due to the degenerate nature of the genetic code, the model also allows synonymous mutations. Thus, the model explores the effects of mutations in both loci, as well as mutations in only one locus. The framework assumes equal mutation rates at both loci, as it proposes a mutation at each locus after every step in the simulation, with 50 replicates of 200 steps of substitution in each simulation. Restricting the mutations to the interface maintains sequence identity above 40%, which has been described previously as the threshold at which protein fold remains similar (Addou et al., 2009; Todd et al., 2001; Wilson et al., 2000).

##### 5.2.2 Implementation of selection

Simulations were carried out using the FoldX suite version 4 (Guerois et al., 2002; Schymkowitz et al., 2005). Starting structures were repaired with the RepairPDB function, mutations were simulated with BuildModel followed by the Optimize function, and estimations of protein stability and binding energy of the complex were done with the Stability and Analyse Complex functions, respectively. Effects of mutations on complex fitness were calculated using methods previously described (Kachroo et al., 2015). The fitness of a complex was calculated from three components based on the stability of protein subunits and the binding energy of the complex using equation 1:

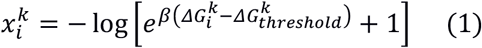

where *i* is the index of the current substitution, *k* is the index of one of the model’s three energetic parameters (stability of subunit A, stability of subunit B, or binding energy of the complex), 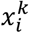 is the fitness component of the *k^th^* parameter for the *I^th^* substitution, *β* is a parameter that determines the smoothness of the fitness curve, 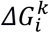 is the free energy value of the *k^th^* free energy parameter (stability of subunit A, stability of subunit B, or binding energy of the complex) for the *i^th^* substitution, and 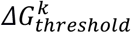 is a threshold around which the fitness component starts to decrease. The total fitness of the complex after the *i^th^* mutation was calculated as the sum of the three computed values for 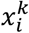, as shown in equation 2:

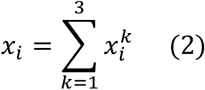

The fitness values of complexes were then used to calculate the probability of fixation (pfix) or rejection of the substitutions using the Metropolis criterion, as in equation 3:

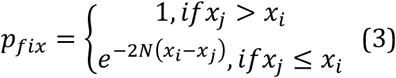

where *p_fix_* is the probability of fixation, *x_i_* is the total fitness value for the complex after *i* substitutions; *x_j_* is the total fitness value for the complex after *j* substitutions, with *j* = *i* + 1; and *N* is the population size, which influences the efficiency of selection.

Different selection scenarios were examined depending on the complexes whose binding energy and subunit stabilities were under selection: neutral evolution (no selection applied on subunit stability and on the binding energy of the complex), selection on one homodimer, selection on the two homodimers, and selection on the heterodimer. *β* was set to 10, *N* was set to 1000 and the 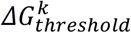 were set to 99.9% of the starting values for each complex, following the parameters described in (Kachroo et al., 2015). For the simulations with neutral evolution, *β* was set to 0. For simulations with other combinations of parameters, we varied *β* and *N*, one at a time, with *β* taking values of 1 and 20 and *N* taking values of 100 and 10000. The simulations with 500 substitutions were carried out with *β* set to 10, and *N* set to 1000.

##### 5.2.3 Analyses of simulations

The results from the simulations were then analyzed by distinguishing mutational steps with only one non-synonymous mutation (single mutants, between 29% and 34% of the steps in the simulations) from steps with two non-synonymous mutations (double mutants, between 61% and 68% of the steps). The global data was used to follow the evolution of binding energies of the complexes over time, which are shown in Figure 4. The effects of mutations in HM and HET were compared using the single mutants (Figure 5-figure supplement 2). The double mutants were used to analyze epistatic and pleiotropic effects (Figure 5, Figure 5-figure supplement 1) and to compare the rates of mutation fixation based on their effects on the HMs (Figure 5–figure supplement 3).

## Author contributions

CRL, AM and AFC designed this study. AM, AKD, IGA, DA, SA, CE and DEY performed the experiments. AFC performed the *in silico* evolution experiments and the analysis of protein structures. AM, AFC, HAJ and CRL analysed the results. CRL and NY supervised the research. AM, AFC and CRL wrote the manuscript with input from all authors.

## Acknowledgements

This work was supported by Canadian Institutes of Health Research grants 299432, 324265 and 387697 to CRL. AM was supported by a FRQS postdoctoral scholarship. AFC was supported by fellowships from PROTEO, MITACS, and Université Laval, as well as joint funding from MEES and AMEXCID. SA was supported by an NSERC undergraduate scholarship. CRL holds the Canada Research Chair in Evolutionary Cells and Systems Biology. We thank SW Michnick for sharing data before publication. The authors thank Philippe Després, Johan Hallin and Anna Fijarczyk for comments on the paper, Rohan Dandage for both comments on the paper and assistance on gathering the data for human paralogs, Rong Shi for useful discussions, and Stéphane Larose for assistance on data management.

## Competing interests

The authors have no competing interests to declare.

## Notes

#### Summary of Updates

Several sections of the text and figures have been modified and additional analyses have been performed.

