## Supplementary text and figures for "The role of structural pleiotropy and regulatory evolution in the retention of heteromers of paralogs"

**Supplementary material of Marchant et al.**

**Table of contents:**


| | Page |
|---|---|
| Supplementary text | 4 |
| Comparison of PCA results with previous studies | 4 |
| Table S1: Homomer data | 4 |
| Table S2: Table S1 Headers | 4 |
| Table S3: Summary results per pair of paralogs | 4 |
| Table S4: Table S3 Headers | 4 |
| Table S5: Description of RNAseq samples | 4 |
| Table S6: Number of RNAseq reads per gene | 4 |
| Table S7: Results from GLM on protein interactions versus expression levels | 4 |
| Table S8: Counts of mutation types in the simulations of protein evolution | 5 |
| Table S9: List of paralog pairs | 5 |
| Table S10: Table S9 Headers | 5 |
| Table S11: Oligonucleotides used in this study | 5 |
| Table S12: Culture media used in this study | 5 |
| Table S13: PDB structures used in this study | 5 |
| Table S14: Summary of results for pairs of human paralogs | 5 |
| Table S15: Table S14 headers | 5 |
| Figure 2-figure supplement 1: Comparison of PCA data generated in this study with published data. | 6 |
| Figure 2-figure supplement 2: Intersections of detected HMs (A) and HETs (B) from this study and previously reported HMs and HETs. | 7 |
| Figure 2-figure supplement 3: Association between mRNA abundance and the probability of HM detection by PCA in this study. | 8 |

Figure 2-figure supplement 4: mRNA and protein abundance of singletons and duplicates. 9

Figure 2-figure supplement 5: Interaction motifs and percentage of pairwise amino acid sequence identity between paralogs. 10

Figure 2-figure supplement 6: Conservation of binding interfaces of human paralogs in HM and HET complexes with solved structures. 11

Figure 3-figure supplement 1: Comparison of Pfam domain composition similarity between pairs of paralogs. 12

Figure 3-figure supplement 2: Comparison of functional similarity between HM and HM&HET pairs. 13

Figure 3-figure supplement 3: Comparison of functional similarity between WGDs, considering homeologs and true ohnologs separately. 14

Figure 3-figure supplement 4: Functional similarity between paralogs as a function of their pairwise amino acid sequence identity. 15

Figure 4-figure supplement 1: Percentage of interaction motifs for SSDs, WGDs and the two types of WGDs. 16

Figure 4-figure supplement 2: Effects of selection on one complex on the other complexes hold for six different PDB structures. 17

Figure 4-figure supplement 3: Effect of changes in parameters on the observed evolution trajectories. 18

Figure 4-figure supplement 4: Single mutants have pleiotropic effects for HM and HET. 19

Figure 5-figure supplement 1: Contribution of epistasis to the evolution of HET for six different PDB structures. 20

Figure 5-figure supplement 2: Distribution of effect sizes of mutations on the binding energy (ΔΔG) of HMs and HETs as estimated using FoldX. 21

Figure 5-figure supplement 3: Fixation rates of double mutants during the simulations. 22

Figure 6-figure supplement 1: The loss of HETs may result from regulatory divergence (single cell RNAseq data (Gasch et al. 2017)). 23

Figure 6-figure supplement 2: Interaction motifs and similarity of functions for SSDs (yellow) and WGDs (blue). 24

Figure 6-figure supplement 3: Expression of whole-genome duplicates (WGDs) and consequences on interaction motifs. 25

Figure 6-figure supplement 4: Similarity of regulation between paralogs as a function of their pairwise amino acid sequence identity. 26

Figure 2-figure supplement 7: Plate organization for DHFR PCA experiments. 27

Figure 2-figure supplement 8: Density of colony size converted to z-score. 28

### Supplementary text

#### Comparison of PCA results with previous studies

We performed a screen using PCA based on the DHFR (Tarassov et al. 2008) to test for PPIs among SSDs and WGDs in *Saccharomyces cerevisiae*, specifically testing for self-interactions (HMs) and interactions between paralogs (HETs). The yeast DHFR PCA detects direct and near direct interactions without disturbing endogenous regulation, giving insight into the role of transcriptional regulation in the evolution of PPIs (Tarassov et al. 2008; Rochette et al. 2014; Barshir et al. 2018; Gagnon-Arsenault et al. 2013). PCA is one of the standard binary methods used to measure direct and near-direct PPIs in yeast and mammalian cells (Titeca et al. 2019). PCA's performance compares to other standard methods when proper controls and analyses are performed. It has been used successfully by our group and others in various contexts since its first application (Schlecht et al. 2017; Celaj et al. 2017; Chrétien et al. 2018; Stynen et al. 2018; Lev, Volpe, and Ben-Aroya 2014).

In general, the PCA signal in our study strongly correlates with results from previous PCA experiments (Stynen et al. 2018; Tarassov et al. 2008) and other publicly available data (Figure 2-figure supplement 1). Roughly 75% of the HMs and HETs detected in our PCA experiments were previously reported (Figure 2-figure supplement 2, Tables S3 and S4), suggesting that most of the HMs and HETs that can be detected with the available tools and in standard conditions have been discovered. While 76 HMs and 47 HETs reported in other studies were not detected in our PCA, our experiments detected 44 HMs and 19 HETs not previously reported (Tables S3 and S4).

#### Table S1: Homomer data

Homomers among proteins from singleton and duplicated genes: results from PCA experiments performed in this study and from published studies and databases.

#### Table S2: Table S1 Headers

Detailed description of Table S3 column headers.

#### Table S3: Summary results per pair of paralogs

Homomers and heteromers of paralogs: results from PCA experiments performed in this study and from published studies and databases.

#### Table S4: Table S3 Headers

Detailed description of Table S1 column headers.

#### Table S5: Description of RNAseq samples

Number of reads before and after cleaning for the RNAseq experiments in the DHFR PCA growth conditions.

#### Table S6: Number of RNAseq reads per gene

RNAseq reads mapped on the *S. cerevisiae* reference genome.

#### Table S7: Results from GLM on protein interactions versus expression levels

(**A**) Impact of expression levels (protein abundance and mRNA) and duplication on the detection of homomers. (**B**) Impact of the percentage of pairwise amino acid sequence identity and function similarity (GO, correlation of genetic interaction profiles, phenotype, localization and transcription factor) on the maintenance of interaction between paralogs after duplication of a HM (HM *versus* HM&HET). (**C**) Impact of the percentage of pairwise amino acid sequence identity and of the expression correlation profiles on the maintenance of interaction between paralogs after duplication of HMs (HM *versus* HM&HET).

For three models, we performed a GLM (binomial family) and then a Likelihood Ratio Test to estimate whether the factors significantly improved the model (assessed by the part of residual deviation explained by the factor). The factors tested and their data sources are described on the right side of each table. The following information is shown: estimates of the linear function and its standard error in brackets (Estimate (standard error) columns) for the intercept and factors tested, and proportion of residual deviance (Residual deviance columns). P-values for each test are indicated next to the estimated values and residual deviance: "*"($p < 0.05$); "**" ($p < 0.01$); "***" ($p < 0.001$). The following information is provided from the Summ function of the jtools R package (Long 2019): number of observations (N); Akaike Information Criterion (AIC); Bayesian Information Criterion (BIC) and pseudo regression (Pseudo R2).

**Table S8: Counts of mutation types in the simulations of protein evolution**

Counts of attempted and fixed mutations are shown for each of the complexes used in the simulations. Double mutants refer to steps in which both loci get a nonsynonymous substitution, single mutants refer to steps in which one of the loci gets a nonsynonymous substitution, and non-mutants refer to steps in which both loci get a synonymous substitution.

**Table S9: List of paralog pairs**

List of DHFR PCA strains from the DHFR collection of Tarassov et al., (2008) and from Diss et al., (2017). We also report if strain constructions were validated by PCR and if new DHFR PCA strains were constructed and validated for this study.

**Table S10: Table S9 Headers**

Detailed description of Table S9 column headers.

**Table S11: Oligonucleotides used in this study**

List of oligonucleotides used for construction and verification of DHFR PCA yeast strains and Gateway destination plasmids.

**Table S12: Culture media used in this study**

Description of media composition.

**Table S13: PDB structures used in this study**

List of PDB structures and biological assemblies used for the analyses of the conservation of interface sequences and the simulations of protein complex evolution. Proteins matched to the chains in that particular structure along with the counts of total residues and interface residues are shown.

**Table S14: Summary of results for pairs of human paralogs**

Interaction of homomers and heteromers of human paralogs: data from the merged BioGRID-IntAct dataset and the PDB.

**Table S15: Table S14 headers**

Detailed description of Table S14 column headers.

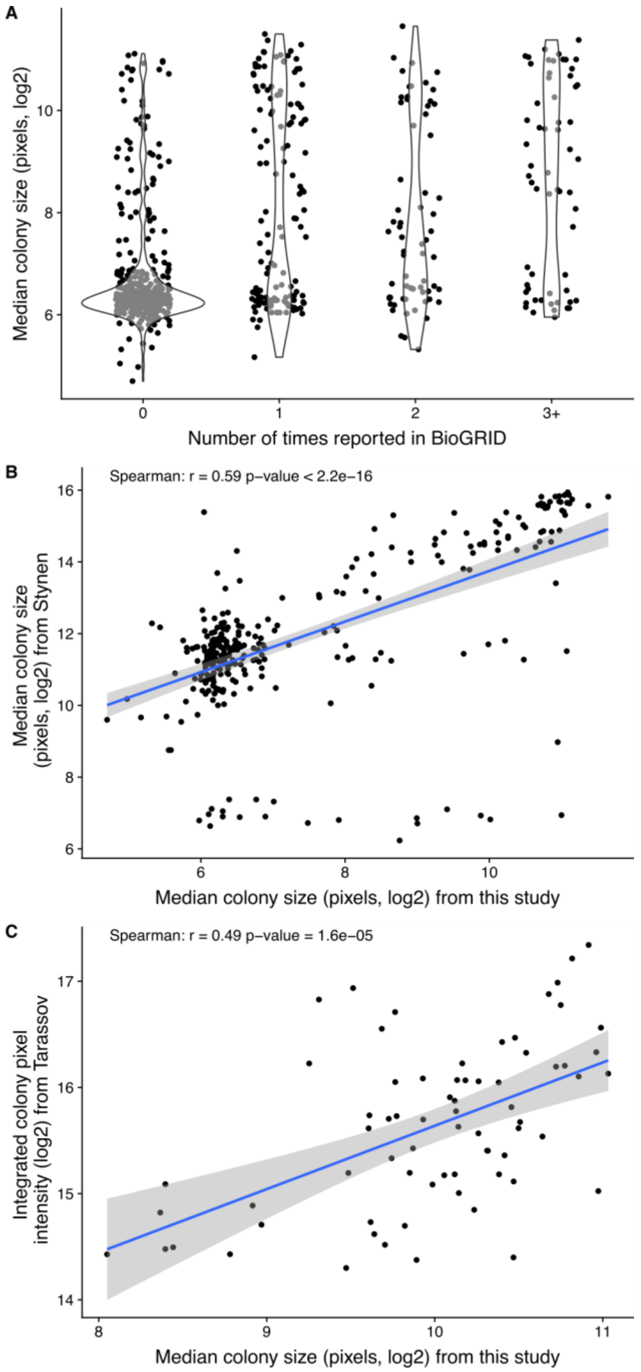


**Figure 2-figure supplement 1: Comparison of PCA data generated in this study with published data.**

(**A**) Colony size (estimated as the integrated pixel intensity) in the PCA experiment as a function of the number of times the corresponding interaction is reported in BioGRID version BIOGRID-3.5.166 (Chatr-Aryamontri et al. 2013, 2017). (**B**) Correlation between colony size of the study of Stynen et al. (Stynen et al. 2018) on homomers and of the PCA experiment performed in this study. (**C**) Correlation between colony size of Tarassov et al. (Tarassov et al. 2008) and of the PCA experiment performed in this study.

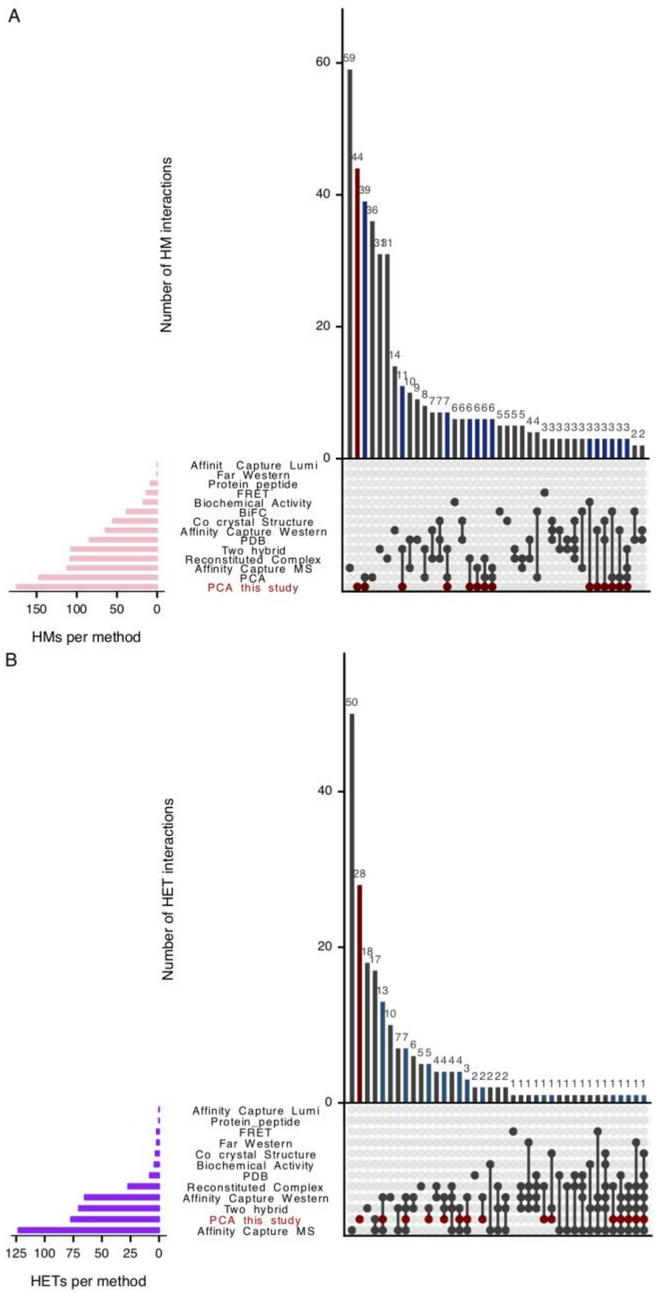


**Figure 2-figure supplement 2: Intersections of detected HMs (A) and HETs (B) from this study and previously reported HMs and HETs.**

We considered HMs and HETs reported in crystal structures from the Protein Data Bank on September 21st, 2017 (Berman et al. 2000) and by PCA based on fluorescent proteins (BiFC) (Kim et al. 2019). We also include HMs and HETs reported in BioGRID (BIOGRID-3.5.166 (Chatr-Aryamontri et al. 2013, 2017) with these methods: Affinity Capture-MS, Affinity Capture-Western, Reconstituted Complex, Two-hybrid, Biochemical Activity, Co-crystal Structure, Far Western, FRET, Protein-peptide, Affinity Capture-Luminescence and PCA. We added data from Stynen et al., (2018) to the BioGRID PCA data. Results of the PCA experiments from this study are highlighted in red. Turquoise-blue bars show HMs and HETs detected in this study and previously observed. The intersections were computed and plotted using the R package UpSetR (Lex et al. 2014).

**Figure 2-figure supplement 3: Association between mRNA abundance and the probability of HM detection by PCA in this study.**

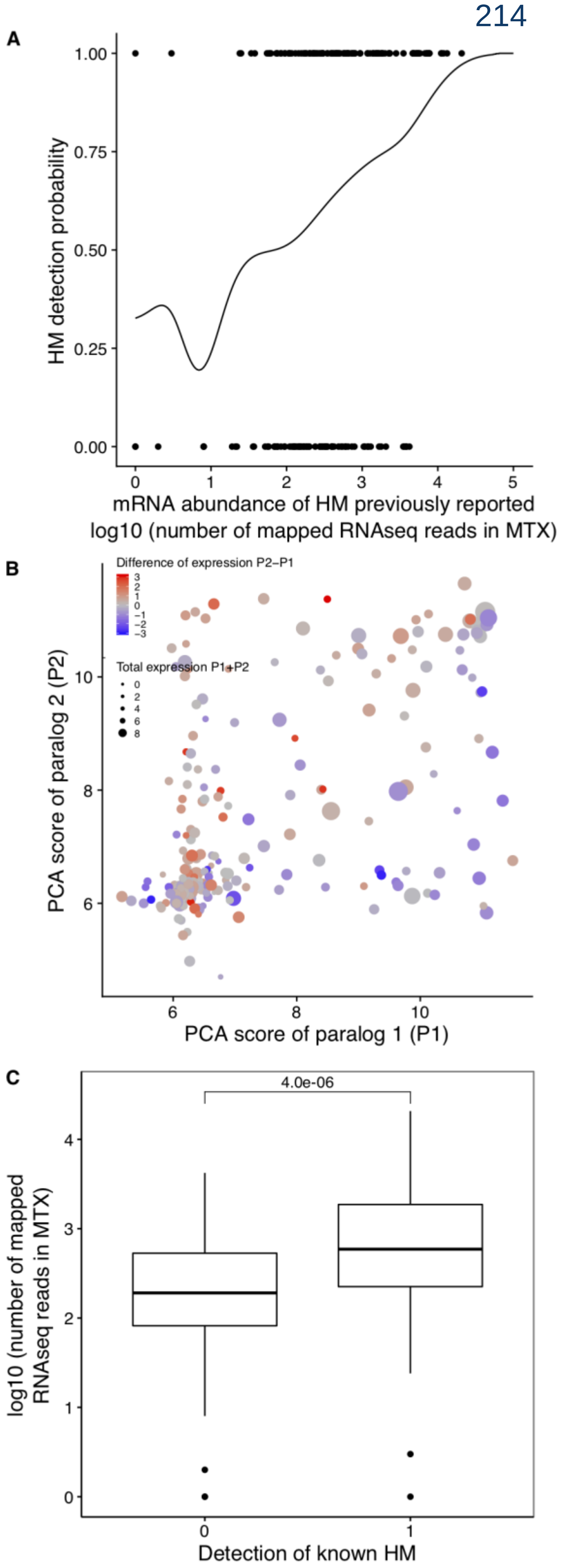


(**A**) The probability that PCA detects a HM is correlated with expression level, as estimated by RNAseq. The plot shows the detection probability of HMs as a function of mRNA abundance for previously reported HMs. Kernel regression of the HM detection (1 for detected, 0 for not detected) on the number of mapped reads per gene ($log_{10}$). (**B**) Difference in HM formation between paralogs results in part from their differential mRNA abundance. The PCA score of paralog 1 (P1) is compared to the PCA score of paralog 2 (P2). PCA scores are median colony sizes from the PCA experiments performed in this study. The total mRNA abundance of paralogs is shown by the size of the points and the difference of expression levels is represented by a color gradient (red for overexpression of P2 compared to P1 and blue overexpression of P1 compared to P2). Red points tend to be above the diagonal, blue points, below the diagonal. (**C**) Comparison of expression levels of previously reported HMs for HMs undetected and detected in the PCA experiments performed in this study. P-value from a Wilcoxon test is shown.

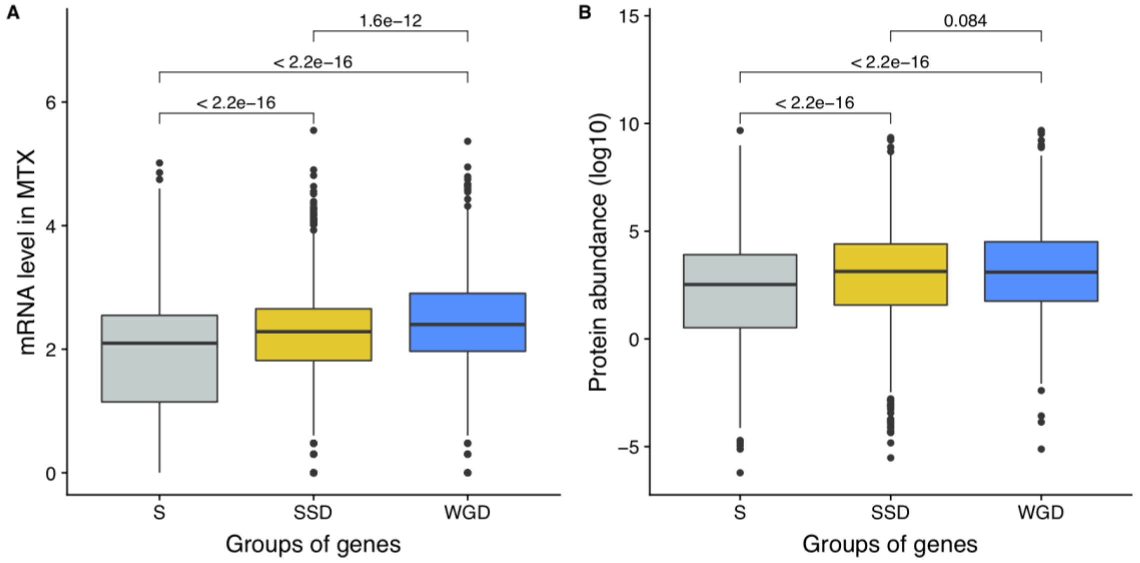


**Figure 2-figure supplement 4: mRNA and protein abundance of singletons and duplicates.**

(**A**) Comparison of mRNA abundance of genes as a function of whether they rare duplicated and of their type of duplication. (**B**) Comparison of the protein abundance as a function of whether they rare duplicated and their type of duplication. (S: singleton, SSD: Small-Scale Duplicates, WGD: Whole-Genome Duplicates). Numbers indicate p-values from Wilcoxon tests.

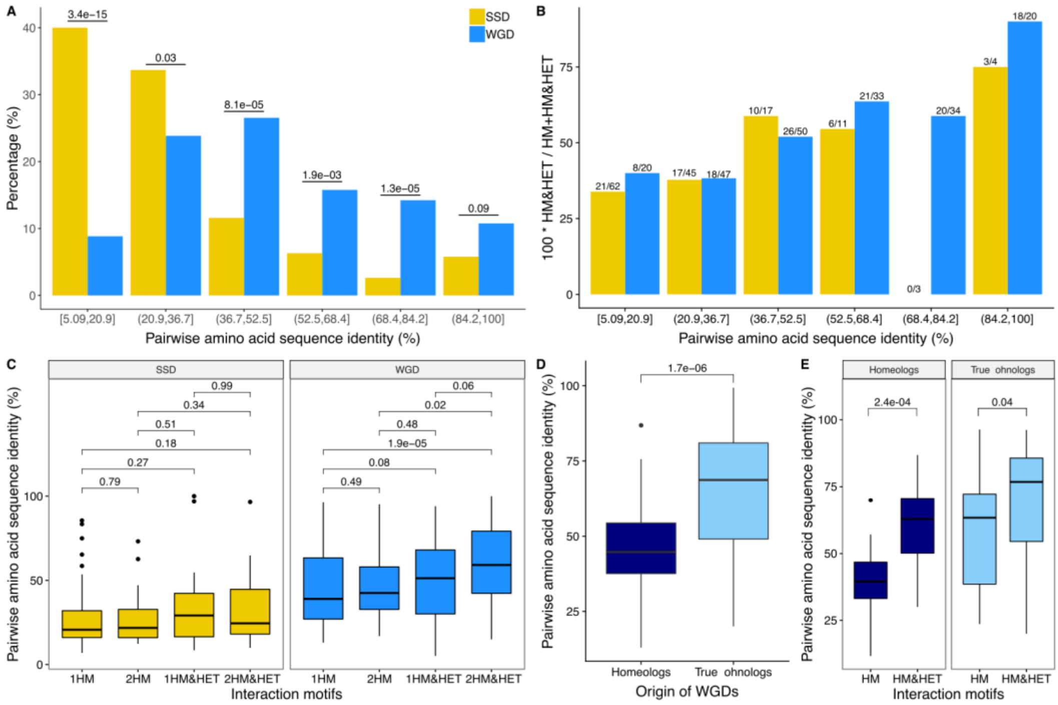


**Figure 2-figure supplement 5: Interaction motifs and percentage of pairwise amino acid sequence identity between paralogs.**

(**A**) Pairs of paralogs were clustered in 6 pairwise amino acid sequence identity groups and the distribution (in percentage) of these groups were compared between SSD and WGD. P-values are from Fisher's exact tests. (**B**) The percentage of paralog pairs forming HM&HETs among the total number of paralog pairs forming HM (HM and HM&HET) is shown as a function of the percentage of pairwise amino acid sequence identity (SSDs in yellow and WGDs in blue). For each group, the number of HM&HET pairs and the total number are indicated above the bars. (**C**) Percentage of pairwise amino acid sequence identity between paralogs for each motif. 1HM: shows one homomer only, 2HM: shows both homomers, 1HM&HET: shows one homomer and the heteromer, and 2HM&HET: shows both homomers and the heteromer. P-values are from Wilcoxon tests. (**D**) The percentage of pairwise amino acid sequence identity among homeologs (dark blue) and true onhologs (light blue). P-value is from a Wilcoxon test. (**E**) Percentage of pairwise amino acid sequence identity between paralogs for HM and HM&HET motifs for homeologs and true ohnologs. P-values are from Wilcoxon tests.

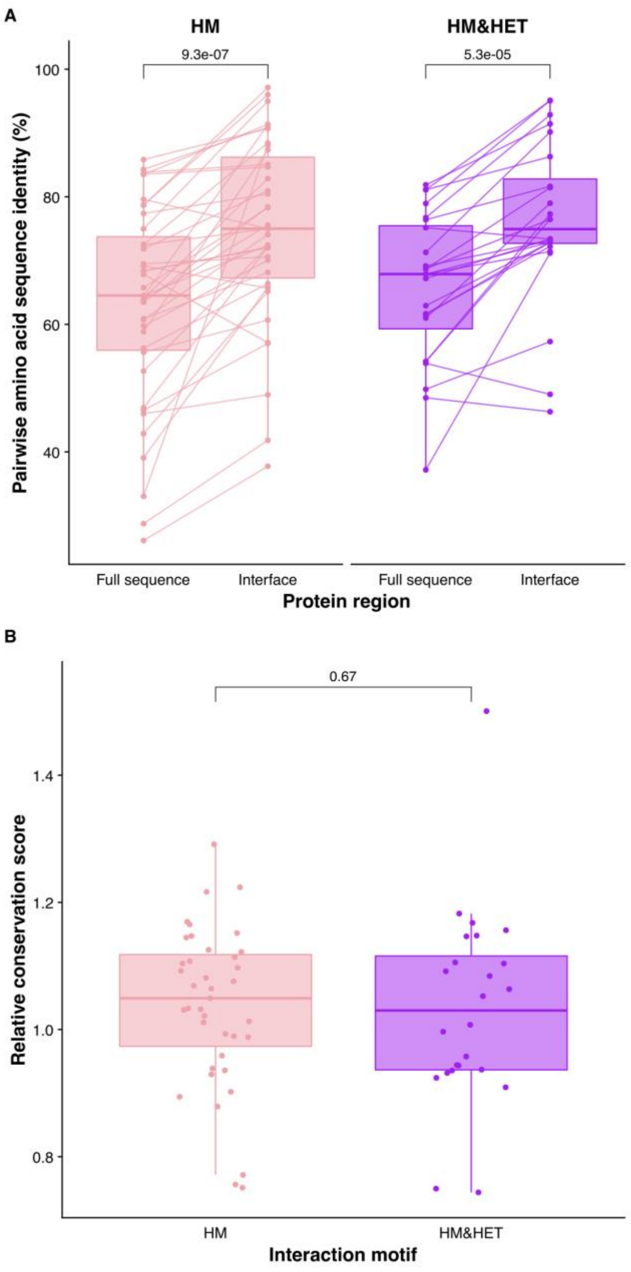


**Figure 2-figure supplement 6: Conservation of binding interfaces of human paralogs in HM and HET complexes with solved structures.**
(**A**) Pairwise amino acid sequence identity for the full sequences of paralogs and their interfaces are shown for the two motifs). P-values from paired Wilcoxon tests are shown. (**B**) Relative conservation scores are shown for the two motifs of paralogs. Relative conservation scores are calculated based on the protein regions solved by crystallography as the percentage of sequence identity at the binding interface divided by the percentage of sequence identity outside the interface. Paralog pairs were classified as HM or HM&HET according to the dataset compiled in Table S14. Homologous interfaces were identified in alignments of the paralogous sequences. Table S13 contains the list of PDB IDs used for these analyses, which include 40 interfaces from homomeric structures for the HM group and 25 interfaces for the HM&HET group (24 homomers and 1 heterodimer of paralogs). P-value is from a Wilcoxon test.

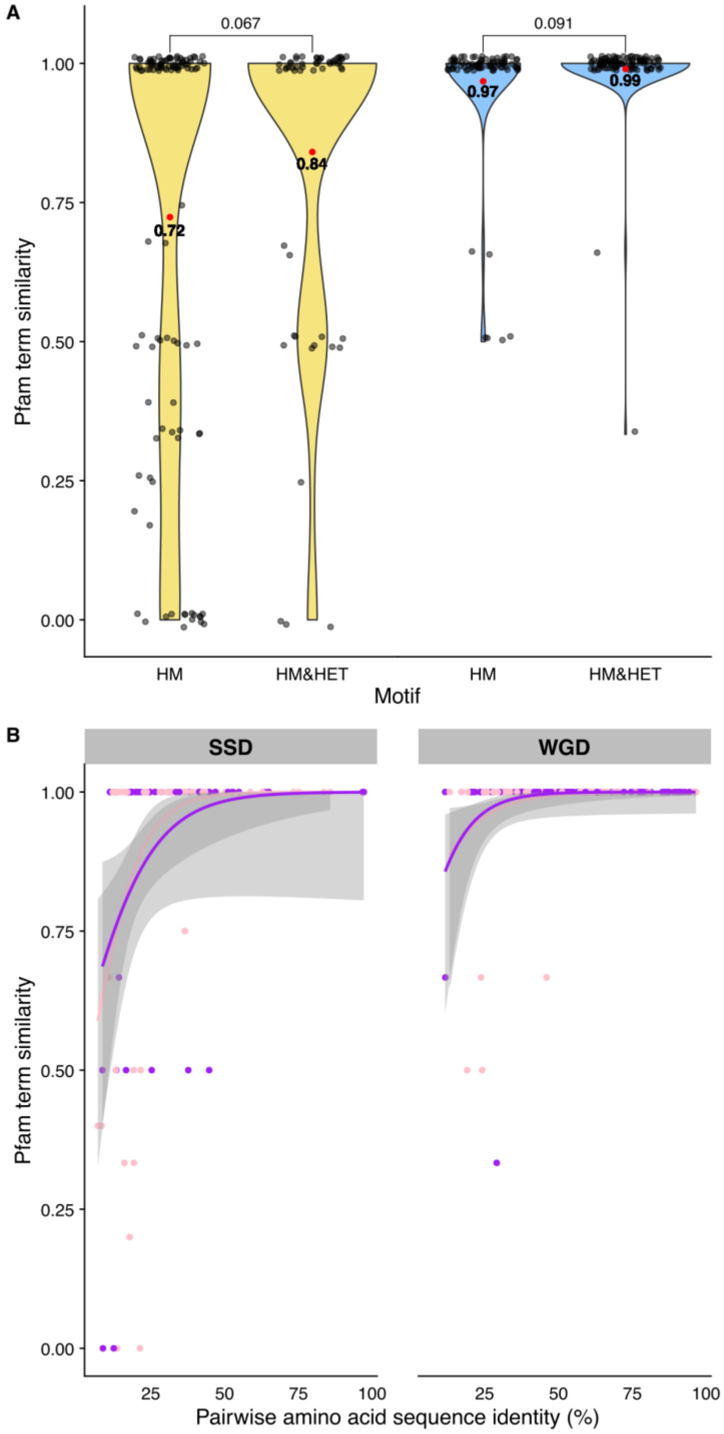


**Figure 3-figure supplement 1: Comparison of Pfam domain composition similarity between pairs of paralogs.**

(**A**) Pfam domain composition similarity (Jaccard's index) between SSDs (yellow) and WGDs (blue) for each interaction motif (HM or HM&HET). (**B**) Pfam domain composition similarity as a function of pairwise amino acid sequence identity for HM motifs (pink) and HM&HET motifs (purple). Regression lines were smoothed using the GLM function with the quasibinomial family.

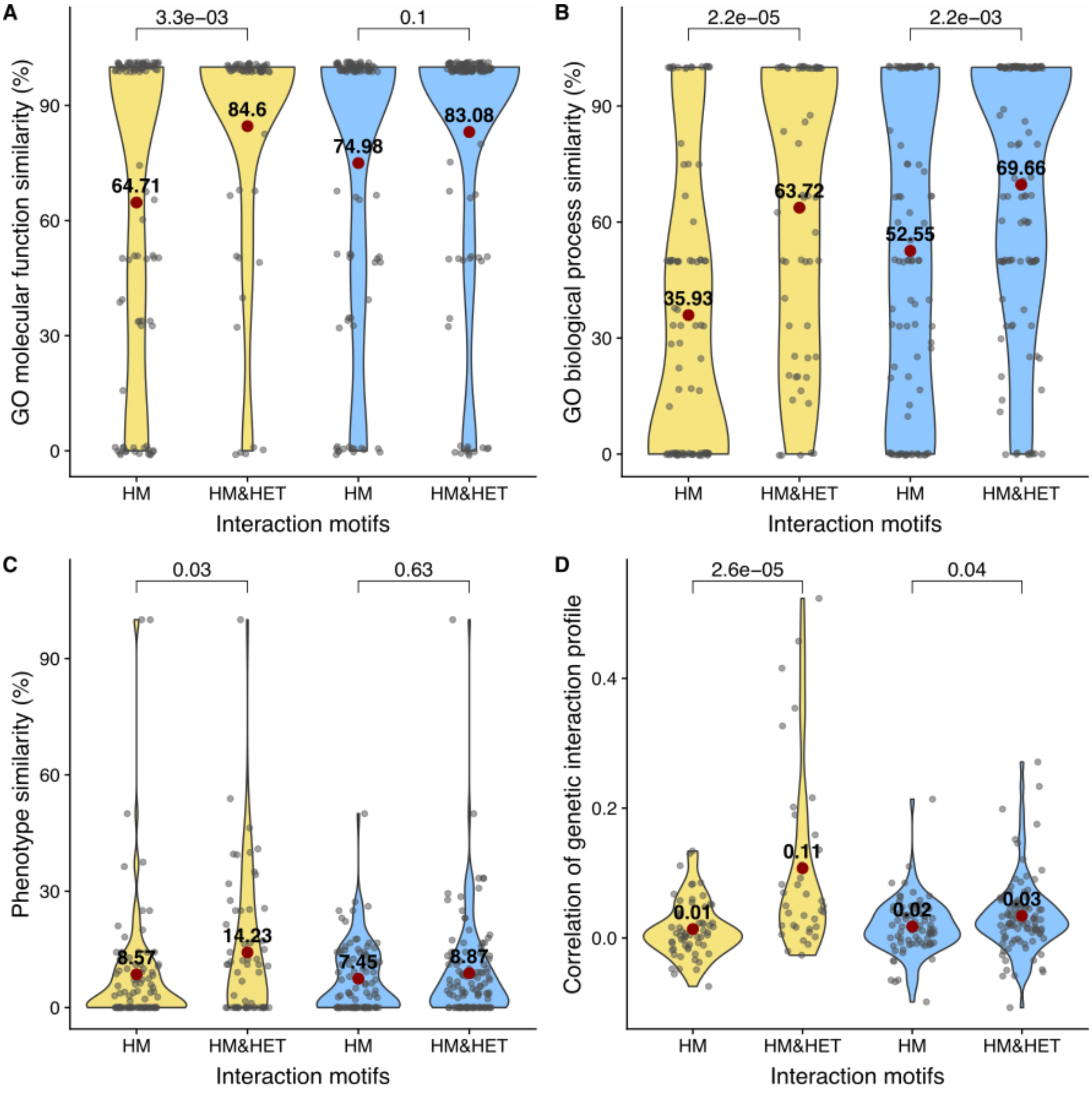


**Figure 3-figure supplement 2: Comparison of functional similarity between HM and HM&HET pairs.**

The similarity of function (100% * Jaccard's index) between SSDs (yellow) and WGDs (blue) was estimated using GO terms for (**A**) molecular functions and for (**B**) biological processes. The similarity of function was also estimated using (**C**) growth phenotypes and (**D**) the correlation of genetic interaction profiles. P-values are from Wilcoxon tests.

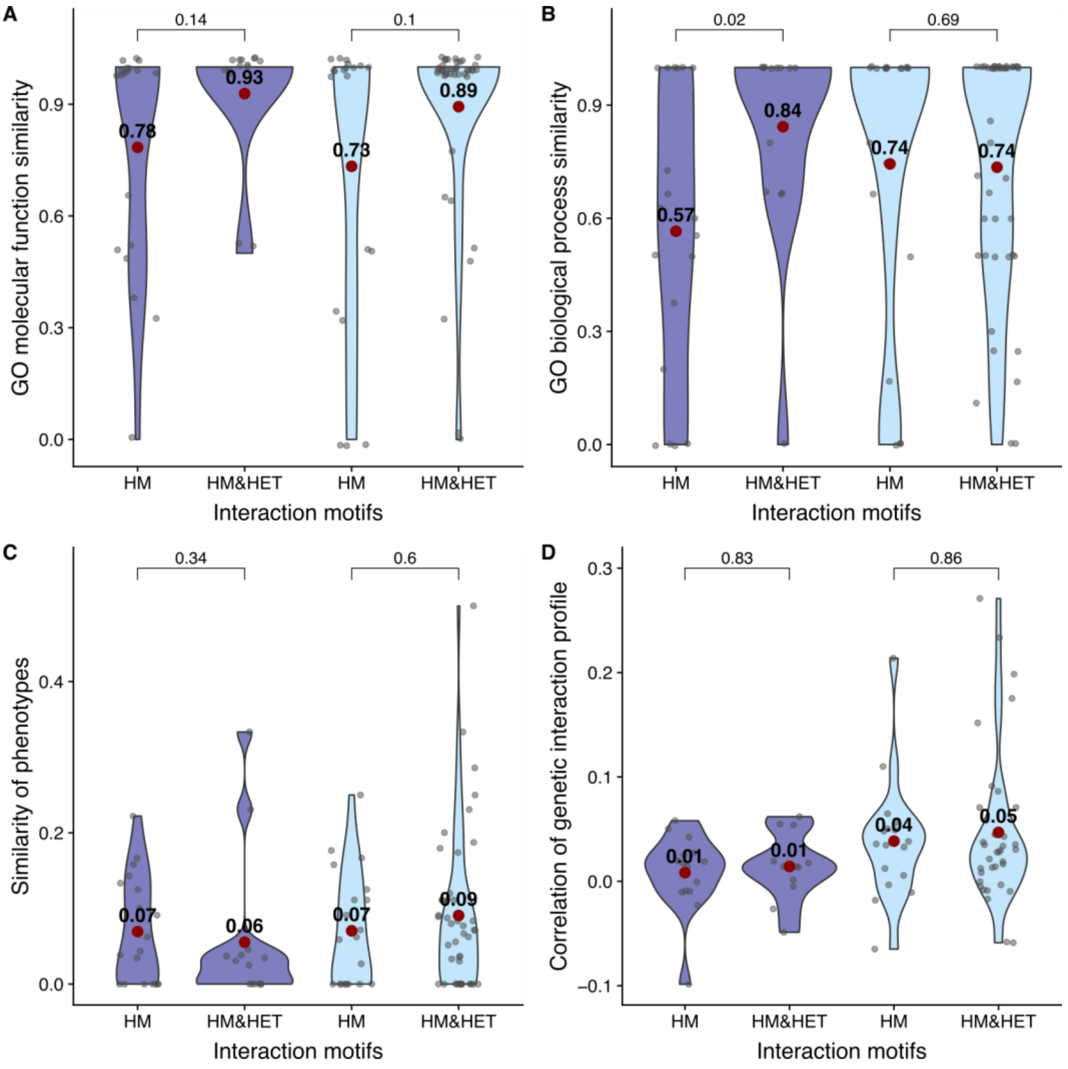


**Figure 3-figure supplement 3: Comparison of functional similarity between WGDs, considering homeologs and true ohnologs separately.**

The similarity of function (100% * Jaccard's index) between homeologs (dark blue) and true ohnologs (light blue) was estimated using GO terms for (**A**) molecular functions and for (**B**) biological processes. The similarity of functions was also estimated using (**C**) growth phenotypes and (**D**) the correlation of genetic interaction profiles. P-values are from Wilcoxon tests.

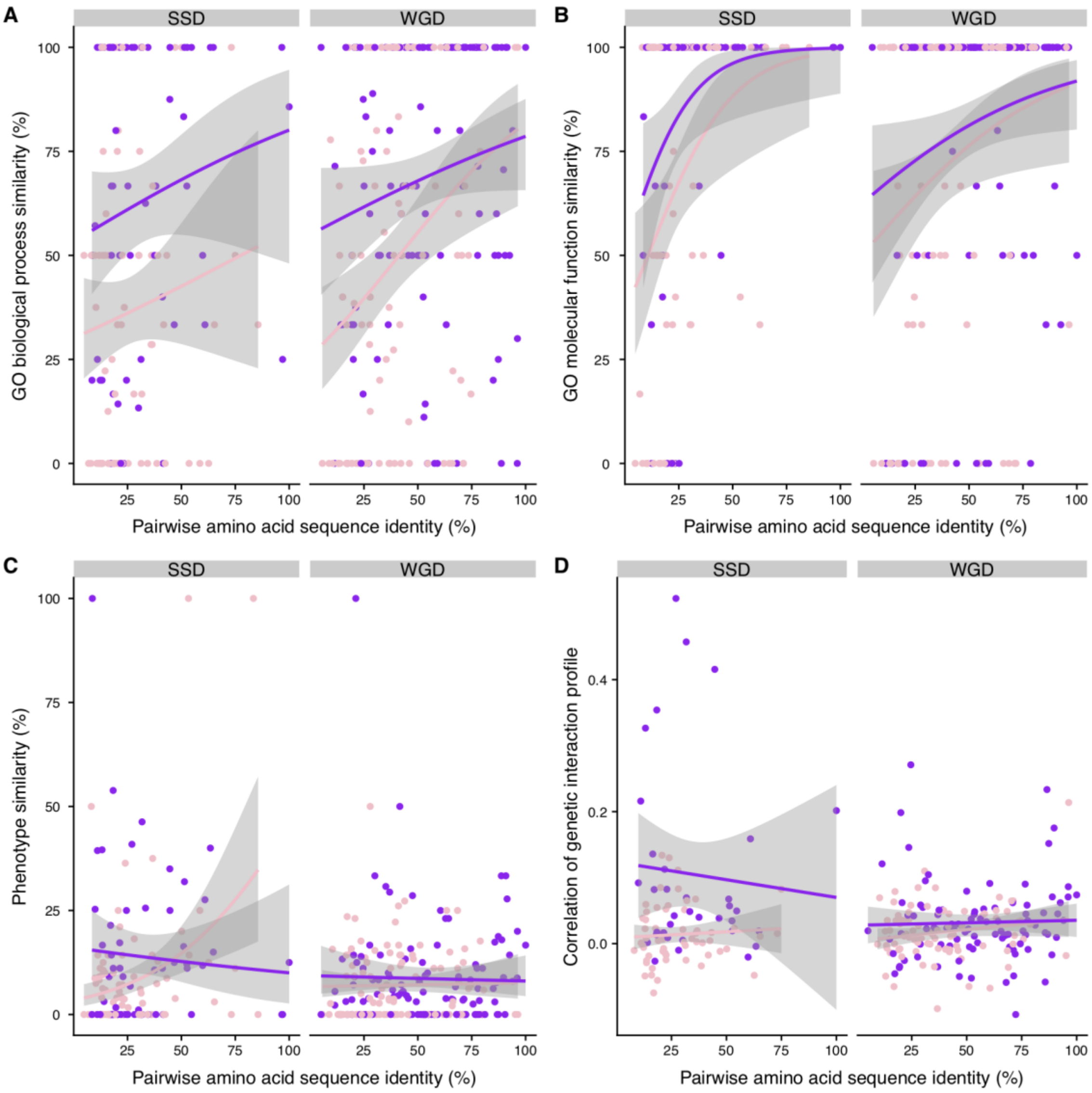


**Figure 3-figure supplement 4: Functional similarity between paralogs as a function of their pairwise amino acid sequence identity.**

The similarity of function (100% * Jaccard's index) between paralogs for HM (pink) and HM&HET (purple) as a function of pairwise amino acid sequence identity for SSDs and WGDs. Similarity of function was estimated using (**A**) molecular functions and (**B**) biological processes GO terms, (**C**) growth phenotypes and (**D**) the correlation of genetic interaction profiles. The regression lines were smoothed using the R geom_smooth function.

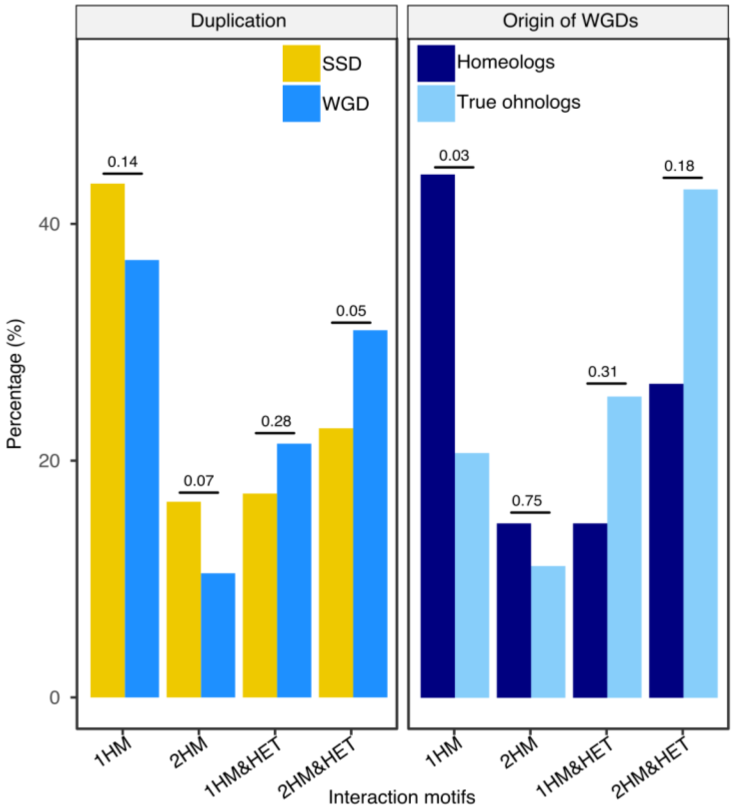


**Figure 4-figure supplement 1: Percentage of interaction motifs for SSDs, WGDs and the two types of WGDs.**

The data is the same as shown in figure 2 but all 4 possible HM and HM&HET motifs are shown. 1HM: shows one homomer only, 2HM: shows both homomers, 1HM&HET: shows one homomer and the heteromer and 2HM&HET: shows both homomers and the heteromer. The percentage of motifs of interaction for SSDs (yellow) and WGDs (blue) (left panel) and for homeologs (dark blue) and true ohnologs (light blue) (right panel). P-values are from Fisher's exact tests.

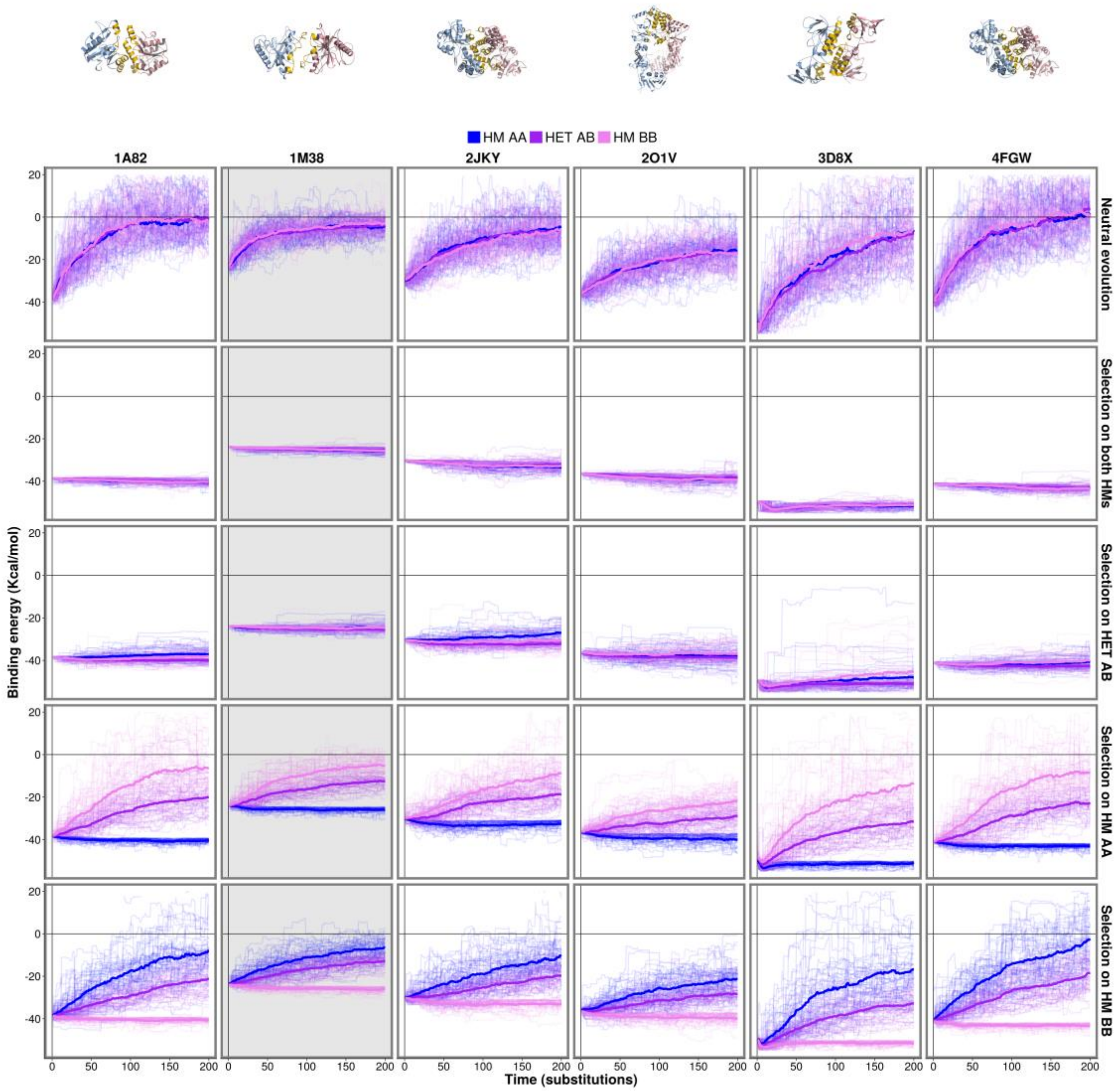


**Figure 4-figure supplement 2: Effects of selection on one complex on the other complexes hold for six different PDB structures.**

The binding energy of six HMs and HETs is followed through time under the same scenarios as shown in Figure 4. Panels shown in Figure 4 are highlighted with a gray background here.

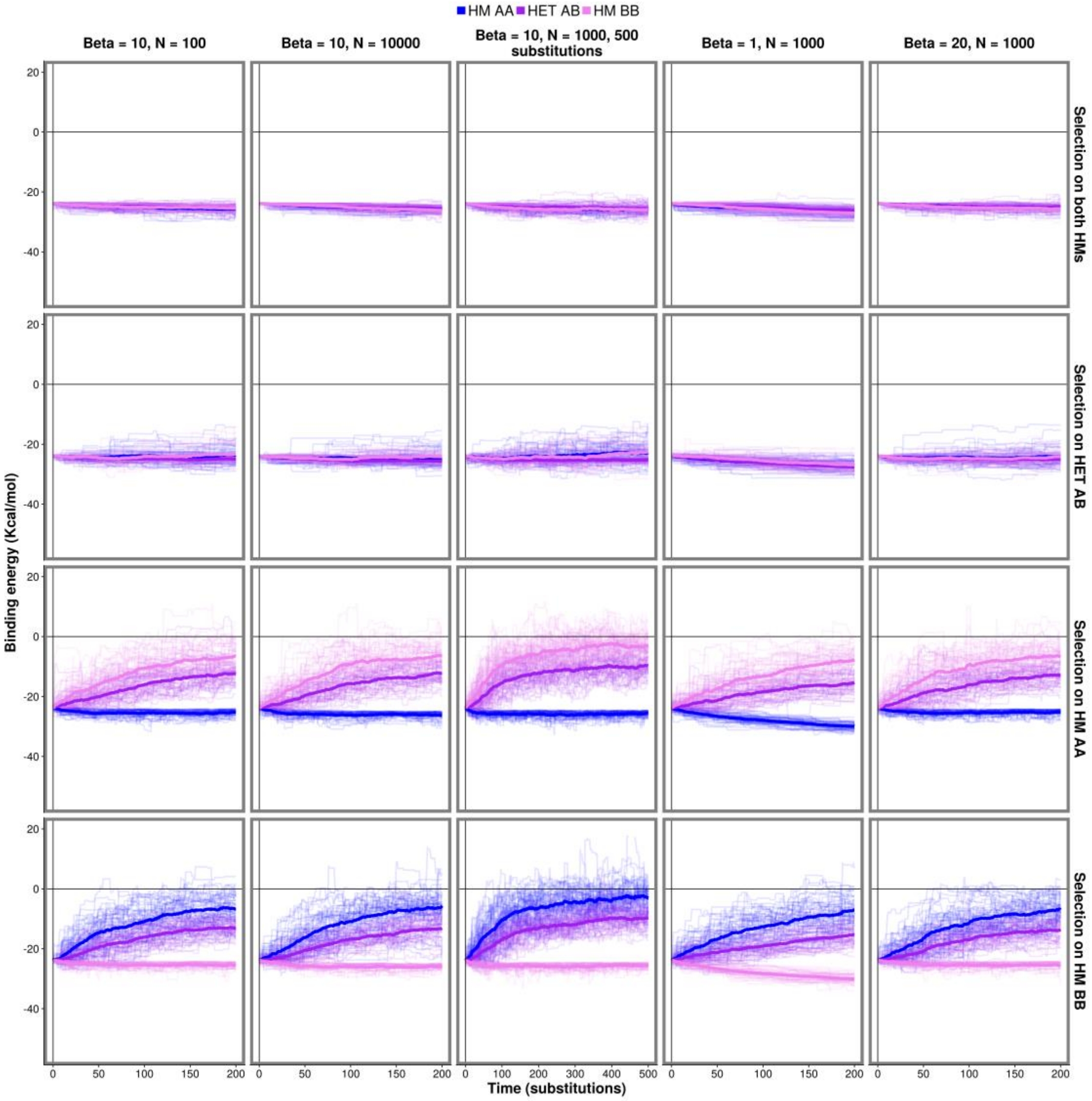


**Figure 4-figure supplement 3: Effect of changes in parameters on the observed evolution trajectories.**

Simulations were run for different combinations of parameters controlling the efficiency of selection ($\beta$ and $N$) and the length of the simulations for PDB structure 1M38.

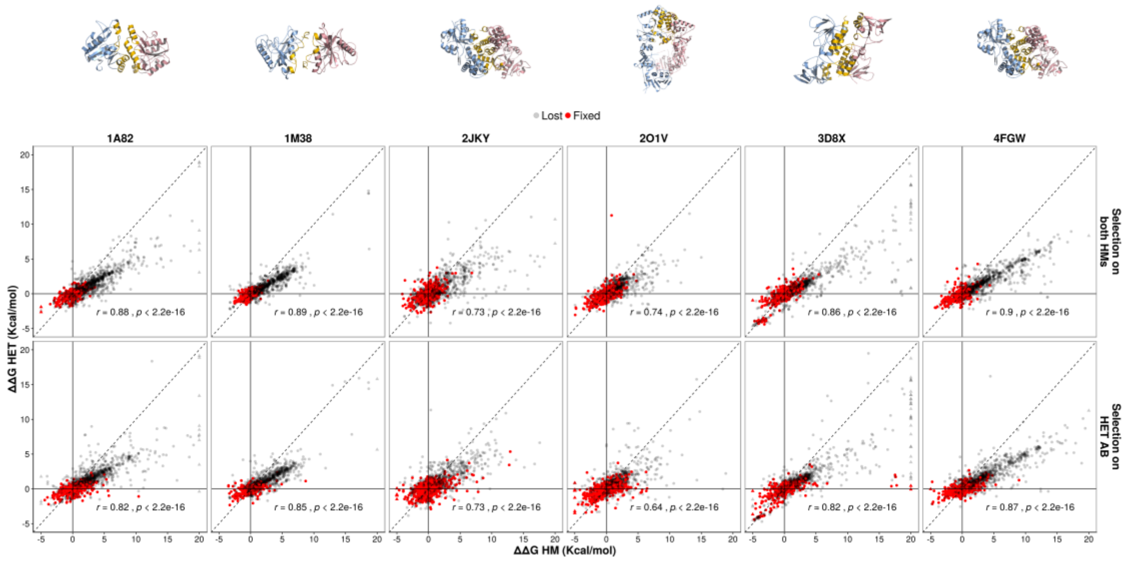


**Figure 4-figure supplement 4: Single mutants have pleiotropic effects for HM and HET.**

The observed effects of sampled single mutants on the HET are compared with their effects on HMs. Pearson correlation coefficients are shown. Parameters used for $\beta$ and $N$ were 10 and 1000, respectively.

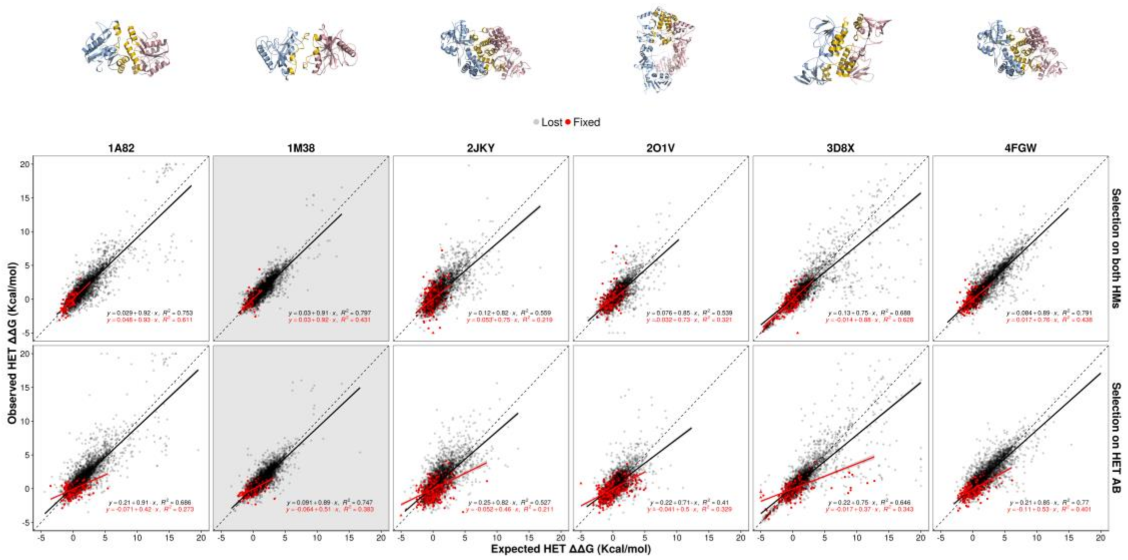


**Figure 5-figure supplement 1: Contribution of epistasis to the evolution of HET for six different PDB structures.**
The observed effects of double mutants on the HET are compared with their expected effects based on the effects on the HMs throughout the simulations. Simulations were run under the same scenarios shown in Figure 5. Panels shown in Figure 5 are highlighted with a gray background. Red points are for mutations that were fixed, grey ones those that were eliminated by selection. The regression equations are shown for fixed and lost mutations separately. Parameters used for $\beta$ and $N$ were 10 and 1000, respectively.

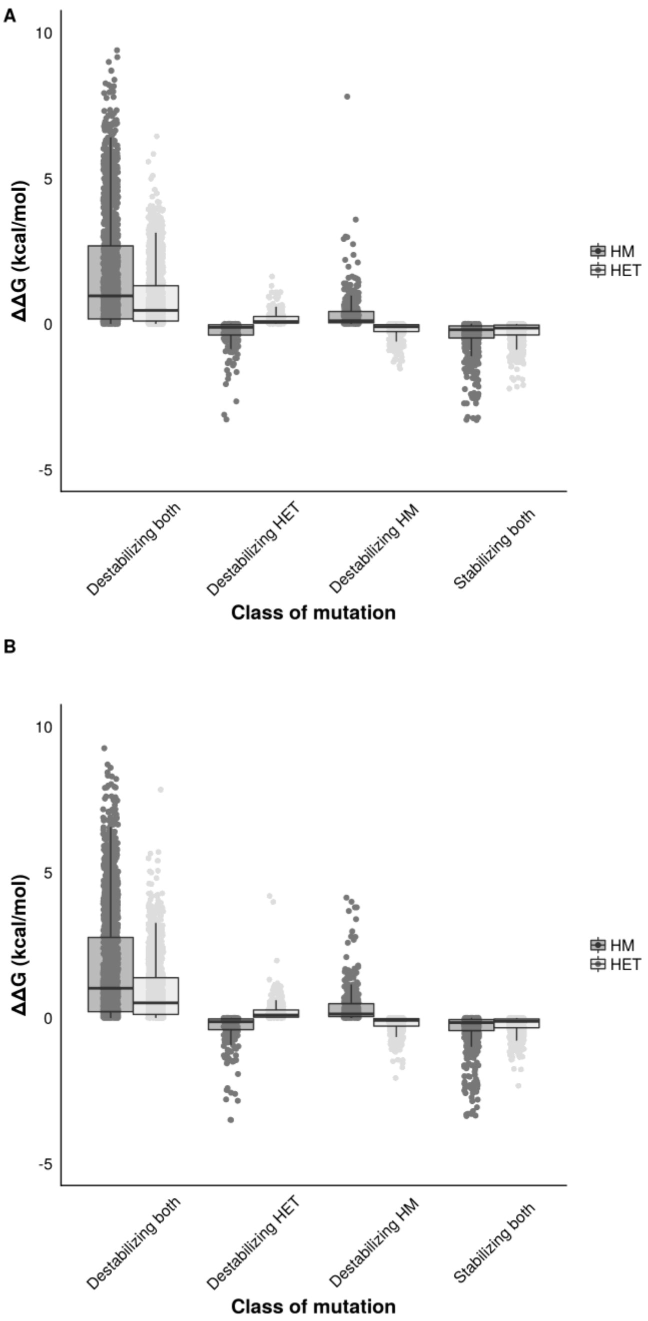


**Figure 5-figure supplement 2: Distribution of effect sizes of mutations on the binding energy (ΔΔG) of HMs and HETs as estimated using FoldX.**
Effects of single mutants on the binding energy of HMs and HETs. Mutants were classified (x-axis) according to their effects on the binding energy of HMs and HETs, depending on whether they stabilize or destabilize both the HM and the HET or they only destabilized one of them. Mutations that destabilize one of the complexes have smaller effect sizes on binding energy than mutations that destabilize or stabilize both. (**A**) Mutations sampled when negatively selecting for the stability of both HMs. (**B**) Mutations sampled when negatively selecting for the stability of the HET. Parameters used for $\beta$ and $N$ were 10 and 1000, respectively.

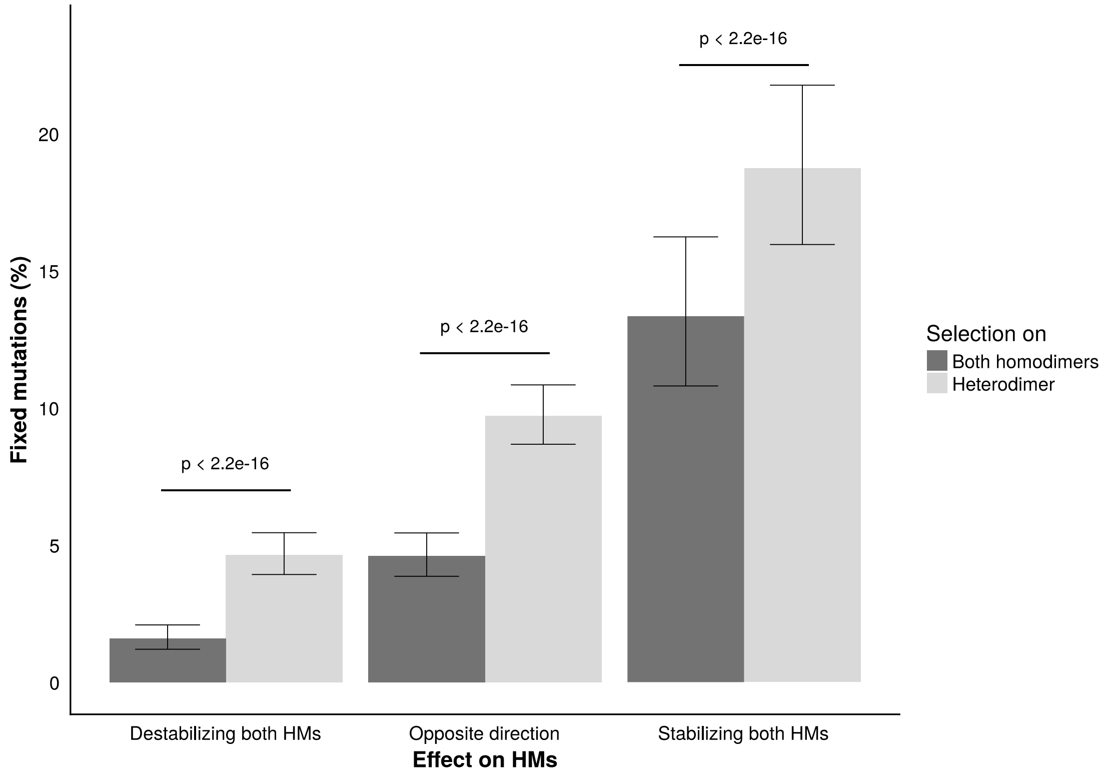


**Figure 5-figure supplement 3: Fixation rates of double mutants during the simulations.**

Fixation rates of double mutants classified based on their effect on the two HMs and the complexes (both HMs or HET) under selection. Clopper-Pearson 95% confidence intervals are shown. P-values were calculated with a two proportion z-test. Parameters used for $\beta$ and $N$ were 10 and 1000, respectively.

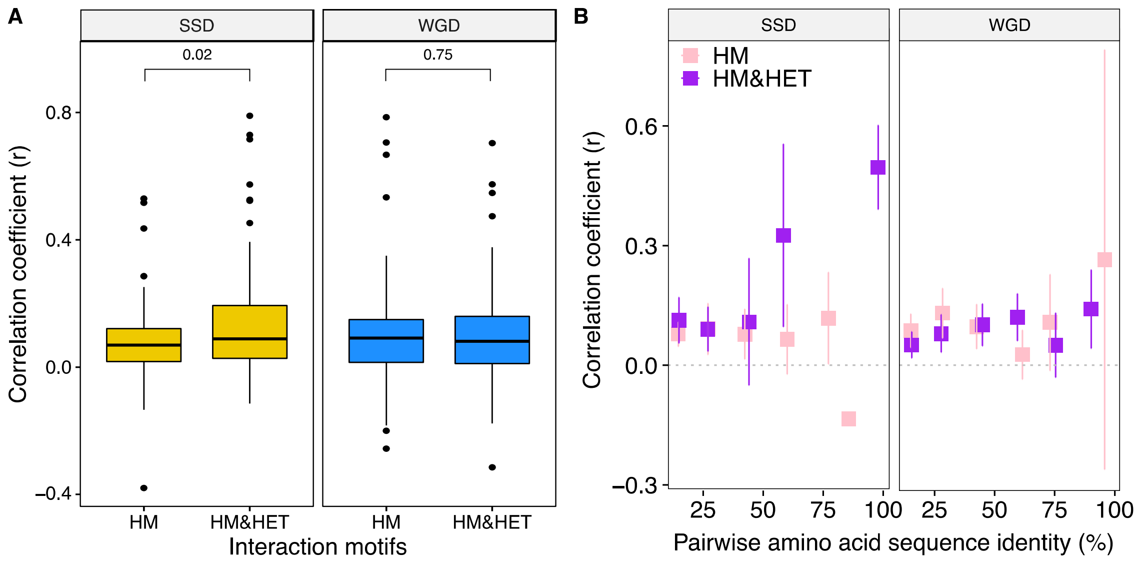


**Figure 6-figure supplement 1: The loss of HETs may result from regulatory divergence (single cell RNAseq data (Gasch et al. 2017)).**

(**A**) Correlation (Spearman r) between the expression profile of paralogs are compared among the different interaction motifs for SSDs (yellow) and WGDs (blue). P-values are from t-tests. (**B**) Correlation of expression profiles between paralogs forming only HM (pink) or HM&HET (purple) as a function of their pairwise amino acid sequence identity.

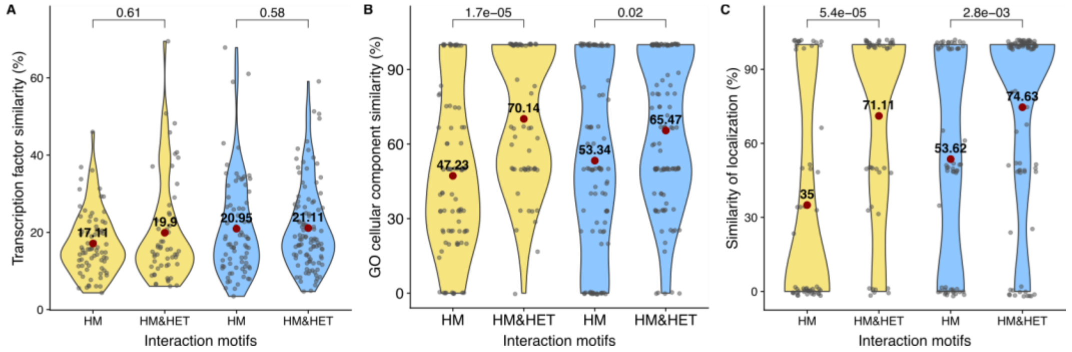


**Figure 6-figure supplement 2: Interaction motifs and similarity of functions for SSDs (yellow) and WGDs (blue).**

The similarity of regulation (100% * Jaccard's index) for (**A**) transcription factor binding sites, (**B**) GO cellular components and (**C**) localization. P-values are from Wilcoxon tests.

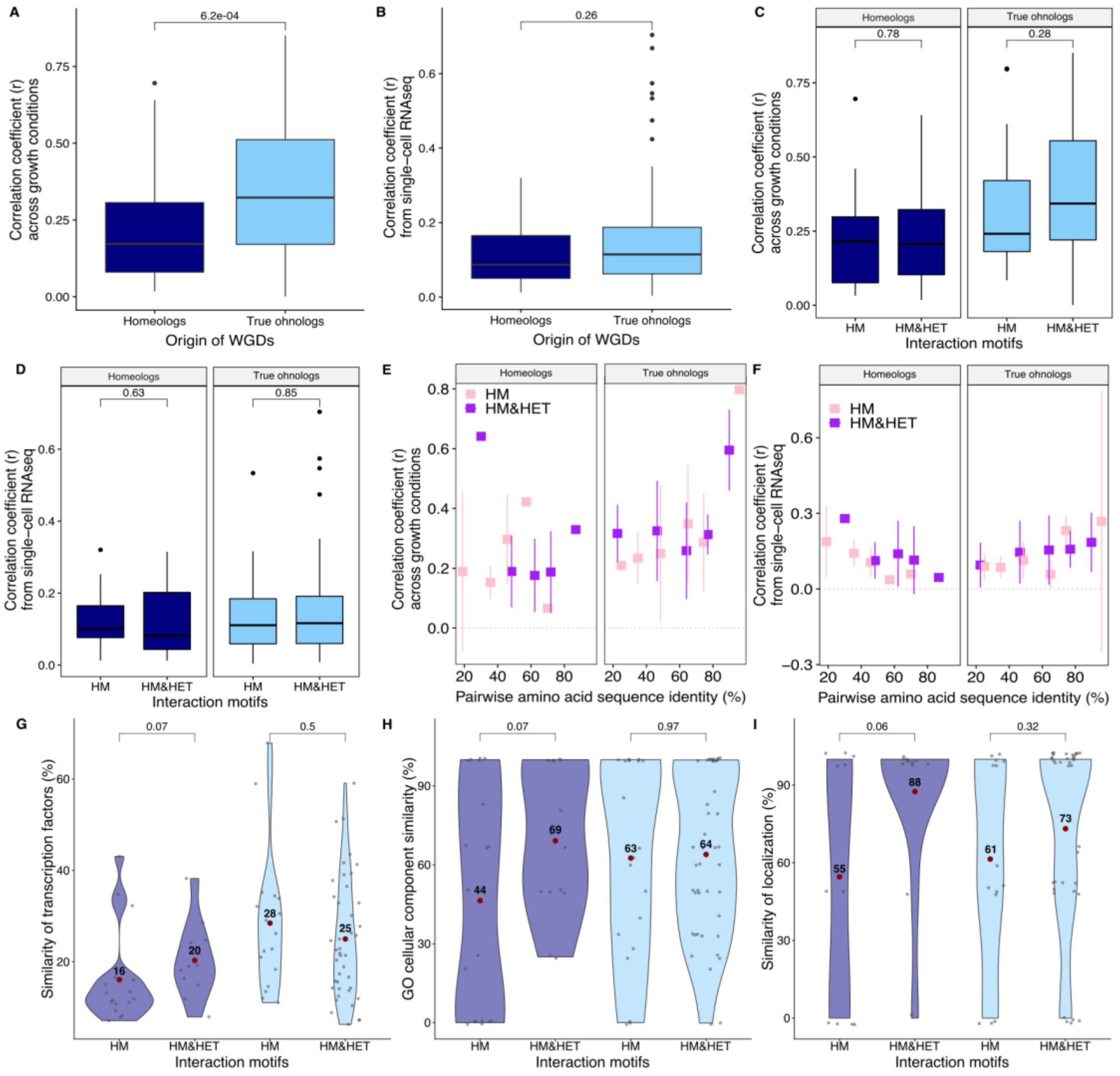


**Figure 6-figure supplement 3: Expression of whole-genome duplicates (WGDs) and consequences on interaction motifs.**

Correlation coefficients (Spearman's r) between the expression profiles of paralogs (**A**) from mRNA relative expression across 1000 growth conditions (Ihmels, Bergmann, and Barkai 2004) and (**B**) from single-cell RNAseq (Gasch et al. 2017) are compared between homeologs and true ohnologs. Correlation coefficients (Spearman's r) (**C**) across growth conditions and (**D**) from single-cell RNAseq data (Gasch et al. 2017) are compared among the different interaction motifs for homeologs and true ohnologs. Correlation coefficients (**E**) across growth conditions and (**F**) from single-cell RNAseq as a function of the percentage of pairwise amino acid sequence identity between paralogs forming only HM or HM&HET. (**G**) Similarity of transcription factor binding sites (100% * Jaccard's index). (**H**) Similarity of GO cellular components. (**I**) Similarity of localization. P-values are from Wilcoxon tests.

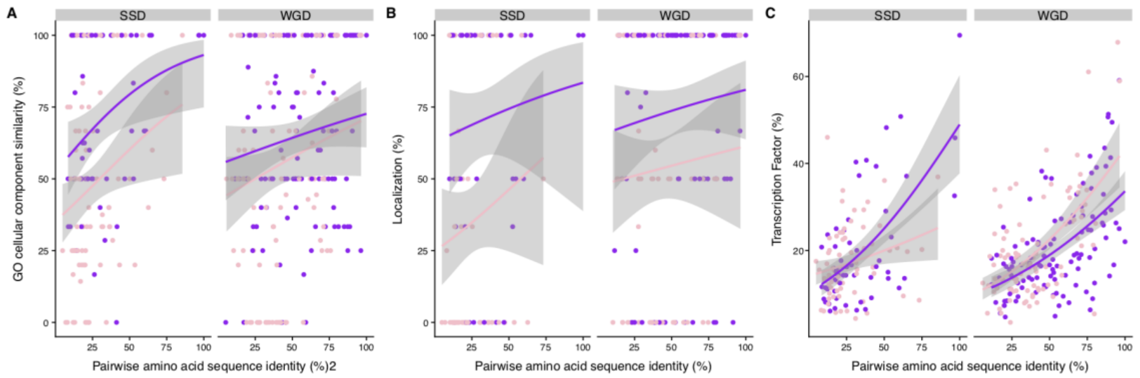


**Figure 6-figure supplement 4: Similarity of regulation between paralogs as a function of their pairwise amino acid sequence identity.**

The similarity of co-expression of HM (pink) and HM&HET (purple) pairs was compared while controlling for pairwise amino acid sequence identity for both SSD and WGD. Similarity of co-expression was estimated using (**A**) cellular component similarity GO term, (**B**) similarity of localization and (**C**) similarity of transcription factor binding sites. The regression lines were smoothed using glm method with quasibinomial family.

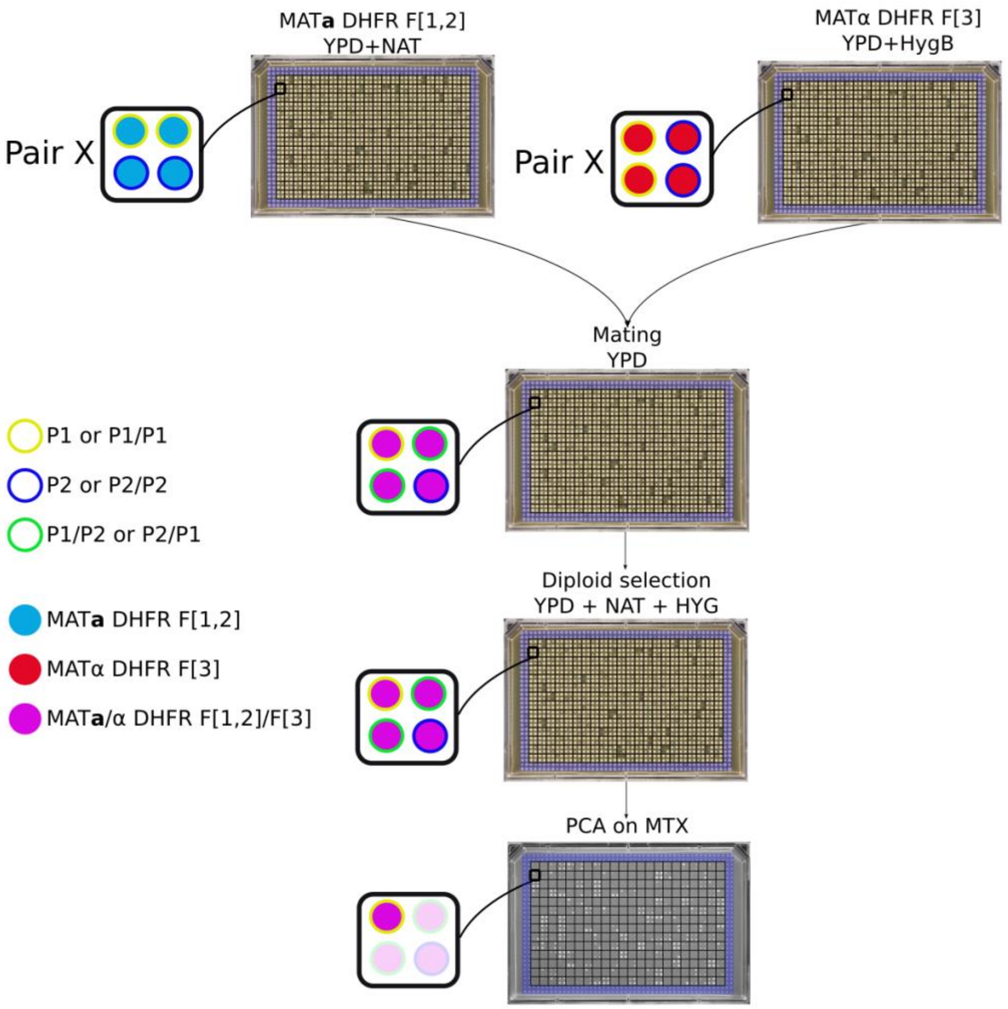


**Figure 2-figure supplement 7: Plate organization for DHFR PCA experiments.** On the haploid arrays (MAT**a** and MATα), each plate has two rows and two columns of control strains at the border (blue lines). Paralogs of a pair are positioned in blocks of four strains. A given pair (example here of pair X) occupies the same position in the MAT**a** and MATα plates. Inside a square, paralogs are positioned horizontally in MAT**a** DHFR F[1,2] plates (P1 are at the top and P2 at the bottom of the square) while they are vertically positioned in MATα DHFR F[3] plates (P1 are at the left and P2 at the right of the square). The two haploid plates were printed on top of each other on a mating plate, generating the following crosses: P1-DHFR F[1,2] / P1-DHFR F[3] at top left, P1-DHFR F[1,2] / P2-DHFR F[3] at top right, P2-DHFR F[1,2] / P1-DHFR F[3] at bottom left and P2-DHFR F[1,2] / P2-DHFR F[3] at bottom right. Two diploid selections and two replications on MTX medium were performed.

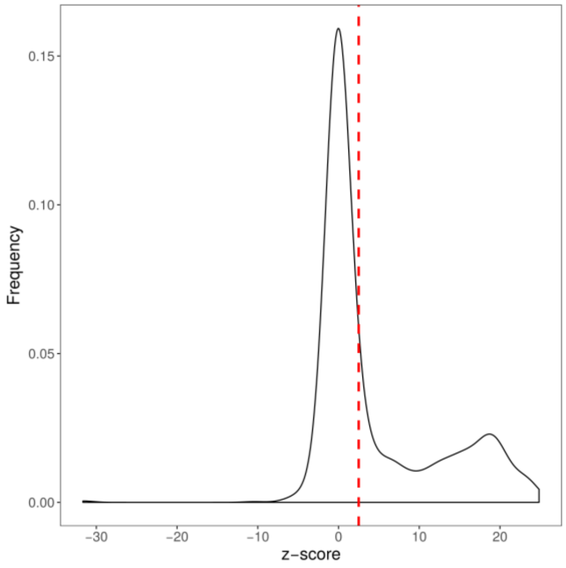


**Figure 2-figure supplement 8: Density of colony size converted to z-score.**
Colony sizes from the PCA experiment of this study were converted to z-score using the mean ($\mu_b$) and standard deviation ($s_{db}$) of the background distribution ($Z_s = (I_s - \mu_b)/s_{db}$)). The density of z-scores is shown in black. A protein-protein interaction was considered as detected if the corresponding z-score was larger than 2.5 (red dashed line).

Schlecht, Ulrich, Zhimin Liu, Jamie R. Blundell, Robert P. St Onge, and Sasha F. Levy.

2017. “A Scalable Double-Barcode Sequencing Platform for Characterization of Dynamic Protein-Protein Interactions.” *Nature Communications* 8 (May): 15586.

Stynen, Bram, Diala Abd-Rabbo, Jacqueline Kowarzyk, Leonor Miller-Fleming, Simran Kaur Aulakh, Philippe Garneau, Markus Ralser, and Stephen W. Michnick. 2018. “Changes of Cell Biochemical States Are Revealed in Protein Homomeric Complex Dynamics.” *Cell*, October. https://doi.org/10.1016/j.cell.2018.09.050.

Tarassov, Kirill, Vincent Messier, Christian R. Landry, Stevo Radinovic, Mercedes M. Serna Molina, Igor Shames, Yelena Malitskaya, Jackie Vogel, Howard Bussey, and Stephen W. Michnick. 2008. “An in Vivo Map of the Yeast Protein Interactome.” *Science* 320 (5882): 1465–70.

Titeca, Kevin, Irma Lemmens, Jan Tavernier, and Sven Eyckerman. 2019. “Discovering Cellular Protein-Protein Interactions: Technological Strategies and Opportunities.” *Mass Spectrometry Reviews* 38 (1): 79–111.
